# Integrator is a genome-wide attenuator of non-productive transcription

**DOI:** 10.1101/2020.07.17.208702

**Authors:** Søren Lykke-Andersen, Kristina Žumer, Ewa Šmidová Molska, Guifen Wu, Jérôme O. Rouvière, Carina Demel, Björn Schwalb, Manfred Schmid, Patrick Cramer, Torben Heick Jensen

## Abstract

Termination of RNA polymerase II (RNAPII) transcription in metazoans relies largely on the Cleavage and Polyadenylation (CPA) and Integrator (INT) complexes originally found to act at the ends of protein-coding and snRNA genes, respectively. Here we monitor CPA- and INT-dependent termination activities genome-wide, including over 8000 previously unannotated transcription units (TUs), that produce unstable RNA. We verify the global activity of CPA, that occurs at pA sites indiscriminately of their positioning relative to the TU promoter. We also identify a global activity of INT, which is, however, largely sequence-independent and restricted to a ~3 kb promoter-proximal region. Our analyses suggest two functions of genome-wide INT activity; it dampens transcriptional output from weak promoters and it provides quality-control of RNAPII complexes, that are unfavorably configured for transcriptional elongation. We suggest that the function of INT in stable snRNA production is an exception from its general cellular role, attenuation of non-productive transcription.

## INTRODUCTION

The RNA output from genomic transcription units (TUs) is determined by transcription initiation and termination. Whereas initiation is critical for the amount of RNA produced, termination at canonical TU ends prevents RNA polymerase (RNAP) from interfering with neighboring TUs and releases the enzyme for new transcription initiation events (Porrua and Libri, 2015; Proudfoot, 2016). Additionally, premature transcription termination at eukaryotic protein-coding loci can radically diminish their production of fulllength transcripts (Arigo et al., 2006; Berg et al., 2012; Elrod et al., 2019; Iasillo et al., 2017; Ntini et al., 2013), which may be subject to regulation (Porrua and Libri, 2015; Proudfoot, 2016). Interrogating mechanisms of transcription termination is therefore central to our understanding of both the integrity of transcriptomes and their expression. A challenge to such inquiries, however, is that transcriptional landscapes are very complex, with RNAP activity occurring pervasively both outside of conventional genic regions and overlapping with other TUs in both sense and antisense orientations (Jensen et al., 2013; Porrua and Libri, 2015; Proudfoot, 2016). A global account of how different termination activities control such ubiquitous transcription is still incomplete.

In metazoans, RNAPII synthesizes all cellular m^7^G-capped RNA and is accountable for the bulk of pervasive transcription. Termination of RNAPII is primarily controlled by two multi-subunit machineries, the Cleavage and Polyadenylation (CPA) complex and the less studied Integrator (INT) complex (Baejen et al., 2017; Guiro and Murphy, 2017; Hsin and Manley, 2012; Kamieniarz-Gdula and Proudfoot, 2019; Proudfoot, 2016; Rosonina et al., 2006). Both termination systems are proposed to rely on the co-transcriptional cleavage of the nascent transcript. This is achieved by the paralogous proteins CPSF3 (CPSF73) and IntS11 (CPSF73L), which are central subunits of CPA and INT, respectively, and the activities of which trigger template release of RNAPII at varying distances downstream of the cleavage site. Transcript cleavage by the CPA complex is instructed by polyadenylation (pA) sites in the nascent RNA, holding at its core a well-defined hexameric AWUAAA consensus element (where W is either A or U) (Proudfoot, 2016). INT-directed cleavage, on the other hand, is suggested to involve a more loosely defined, so-called 3’-box sequence (Baillat and Wagner, 2015; Guiro and Murphy, 2017; Hernandez, 1985), although the generality of this has recently been questioned (Elrod et al., 2019; Tatomer et al., 2019). Nonetheless, transcription termination pathways have traditionally been assigned to discrete classes of transcripts (RNA biotypes); i.e. the presence of pA sites and 3’-box sequences at the ends of protein-coding and snRNA-genes, respectively, has established these RNA biotypes as corresponding archetypical CPA and INT substrates.

Notwithstanding these differences in requirements for CPA and INT functions, these two complexes for 3’-end processing and transcription termination also share characteristics. For instance, they both appear to operate when RNAPII progress is considerably reduced. RNAPII passage across a pA site induces its slow-down, presumably due to structural rearrangements within the transcription complex (Cortazar et al., 2019; Zhang et al., 2015). RNAPII release from the DNA template may be aided by the 5’-3’ degradation of the CPA-produced uncapped transcript that emanates from the RNAPII exit channel (Fong et al., 2015; Kim et al., 2004; West et al., 2004). Likewise, RNAPII decelerates downstream of 3’-box sequences (Cuello et al., 1999; Fong et al., 2015; O’Reilly et al., 2014) in preparation for its termination, and it was recently demonstrated that INT-directed cleavage can occur at a subset of promoter-proximally stalled RNAPII in both drosophila S2 and human HeLa cells (Elrod et al., 2019; Tatomer et al., 2019). This latter activity releases a short transcript and attenuates further transcription elongation to variable degrees.

Given the elusive nature of the 3’-box sequence and provided that pA sites are also used outside of their conventional context at protein-coding gene ends, it is hard to reconcile a model where CPA and INT complexes would act in strict RNA biotype-specific manners (Berg et al., 2012; Kamieniarz-Gdula et al., 2019; Ntini et al., 2013). Indeed, ample ‘cross-reactivity’ of the two termination pathways can occur. For example, INT activity has been reported in both early and late transcriptional stages of signal responsive protein-coding genes (Elrod et al., 2019; Gardini et al., 2014; Lai et al., 2015; Skaar et al., 2015; Stadelmayer et al., 2014) and, conversely, in the absence of INT, transcription termination at snRNA genes can be facilitated by CPA complex components at downstream pA sites (Yamamoto et al., 2014). Moreover, transcription termination at human enhancer/enhancer-like loci, expressing short and labile enhancer RNAs (eRNAs), has been suggested to depend on both CPA and INT (Andersson et al., 2014b; Lai et al., 2015). Finally, subsets of the sizeable, but rather ill-defined, class of mammalian long non-coding RNA (lncRNA) loci were reported to display both pA site-independent (Schlackow et al., 2017; Wu et al., 2020) and pA site-dependent (Almada et al., 2013; Ntini et al., 2013) RNAPII termination. Adding to this complexity, lncRNAs are generally short-lived due to their efficient targeting by the ribonucleolyic RNA exosome (Schmid and Jensen, 2018), rendering annotation of such loci difficult. Altogether, these observations suggest that an RNA biotype-centric view of transcription termination requirements falls short of describing the full substrate repertoire, and functional portfolios, of CPA and INT.

Here, we inquire where CPA and INT activities operate and cooperate over the human genome. We take an RNA biotype-agnostic approach in comparing deep transient transcriptome sequencing (TTseq) and 3’-end-sequencing (3’-end-seq) profiles of cells individually depleted for the CPA subunit CPSF3 or the INT subunit IntS11. Considering the abundant expression of unstable lncRNA, we annotate the HeLa test cell transcriptome by employing data from both unperturbed cells and from cells depleted for the core RNA exosome component EXOSC3 (RRP40), allowing the inclusion of 8027 previously unannotated TUs. Our analyses reveal a global activity of CPA at promoter-distal and -proximal pA sites. Intriguingly, INT also operates globally and with no RNA biotype-specificity, but in contrast to CPA, its activity is restricted to ~3 kb promoter-proximal regions. We suggest that this activity is exploited by snRNA genes, which appear to have developed stronger consensus, and possibly post-transcriptional, INT cleavage sites for their stable 3’-end processing. Our further observations suggest two unprecedented, and non-exclusive, functions of INT activity in dampening the transcriptional output from weak promoters and in providing a quality-control checkpoint to terminate RNAPII complexes that are unfavorably configured for transcriptional elongation and co-transcriptional RNA processing.

## RESULTS

### Exhaustive *de novo* annotation of HeLa cell TUs

To enable an RNA biotype-agnostic analysis of transcription termination, it is critical to monitor newly synthesized RNA and to establish a robust global definition of TUs and the reciprocal intergenic – i.e. truly non-transcribed – regions. We therefore performed transient transcriptome sequencing (TT-seq; (Schwalb et al., 2016)) and total RNA sequencing (RNA-seq) in HeLa cells to capture both newly synthesized and mature RNA from cells either control (CTRL)-treated or subjected to siRNA-mediated depletion of the RNA exosome component EXOSC3 (EXO) (Figure 1A and S1A). The latter condition was chosen to improve the annotation of the numerous exosome-sensitive transcripts (Schmid and Jensen, 2018). A HeLa cell-specific annotation of TUs was conducted using the genomic state annotation (STAN) tool (Zacher et al., 2014; Zacher et al., 2017) as well as comprehensive information about capped transcript 5’-(Andersson et al., 2014b) and 3’-ends (Wu et al., 2020) from EXO-depleted HeLa cells (Figure 1B-C and S1B-C, see *Materials and Methods* for details) (Michel et al., 2017; Schwalb et al., 2016). Gratifyingly, the resulting annotation encompasses more of our datasets as compared to reference annotations from Gencode and RefSeq (Figure S1D). Moreover, since only active loci were included, the fraction of annotations covered by data roughly doubled compared to the reference annotations (data not shown).

**Figure 1.**
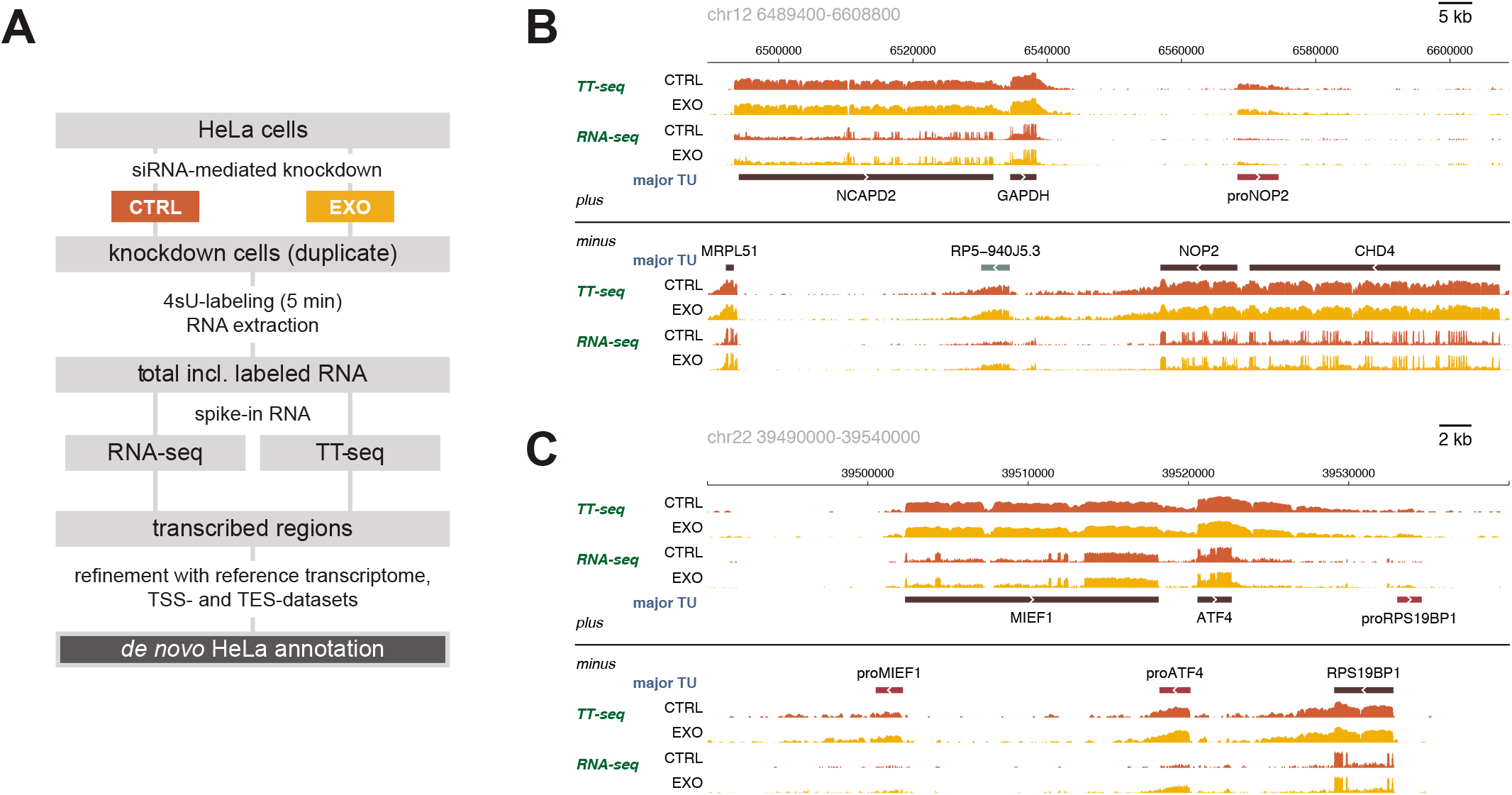
Exhaustive *de novo* annotation of HeLa cell TUs. (A) Flow diagram illustrating the experimental steps underlying annotation of the HeLa cell transcriptome. See text and Figure S1 for details. (B) Genome browser view of chr12:6489400-6608800 illustrating TT-seq and RNA-seq mean coverage signals from control samples – siEGFP- (*red*; CTRL) or siEXOSC3-treated (*orange*; EXO) cells for the plus and minus strand. Proteincoding, antisense as well as previously unannotated TUs (*proNOP2*, a PROMPT, in this view) are displayed in black, gray and red, respectively. (C) As in (B), but for chr22:39490000-39540000. Mean of two replicates are shown for all tracks in (B)-(C). See Figure S1 for additional data related to this figure.

Altogether we defined 22844 TUs, of which 8027 were newly annotated (NA), constituting a substantial size, only exceeded by 11088 protein-coding TUs (Figure S1E). NA TUs were further sub-categorized as ‘enhancer RNA’, based on their overlap with a comprehensive enhancer dataset (Xiong et al., 2018), or ‘promoter upstream transcript (PROMPT)’, ‘intergenic’, ‘intragenic’, ‘natural antisense transcript (NAT)’, ‘overlapping convergent’ or ‘nearby convergent’ TUs according to their genomic locations (Figure S1F-G). Compared to protein-coding genes all of the NA-subtype loci are generally shorter, display lower transcription and steady-state transcript levels and fewer introns (most often none; Figure S1G). Collectively, this division of the HeLa genome into transcribed and non-transcribed regions allowed a comprehensive analysis of transcription termination.

### CPA and INT depletions display common and diverse global transcription termination phenotypes at TU ends

To explore the genome-wide roles of the CPA and INT complexes in transcription termination, we used siRNAs to deplete CPSF3 (CPA) or its paralogue IntS11 (INT) (Figure S1A) and performed TT-seq and RNA-seq (Figure 2A). CTRL- and EXO-depleted cells served as controls. CPA and INT have well-described functions in RNA 3’-end processing and transcription termination at proteincoding and snRNA genes, respectively (Guiro and Murphy, 2017; Proudfoot, 2016)). In agreement, their depletion caused the expected transcription termination defects (see Figure 2B for individual examples and Figure S2A,J for aggregate analyses).

**Figure 2.**
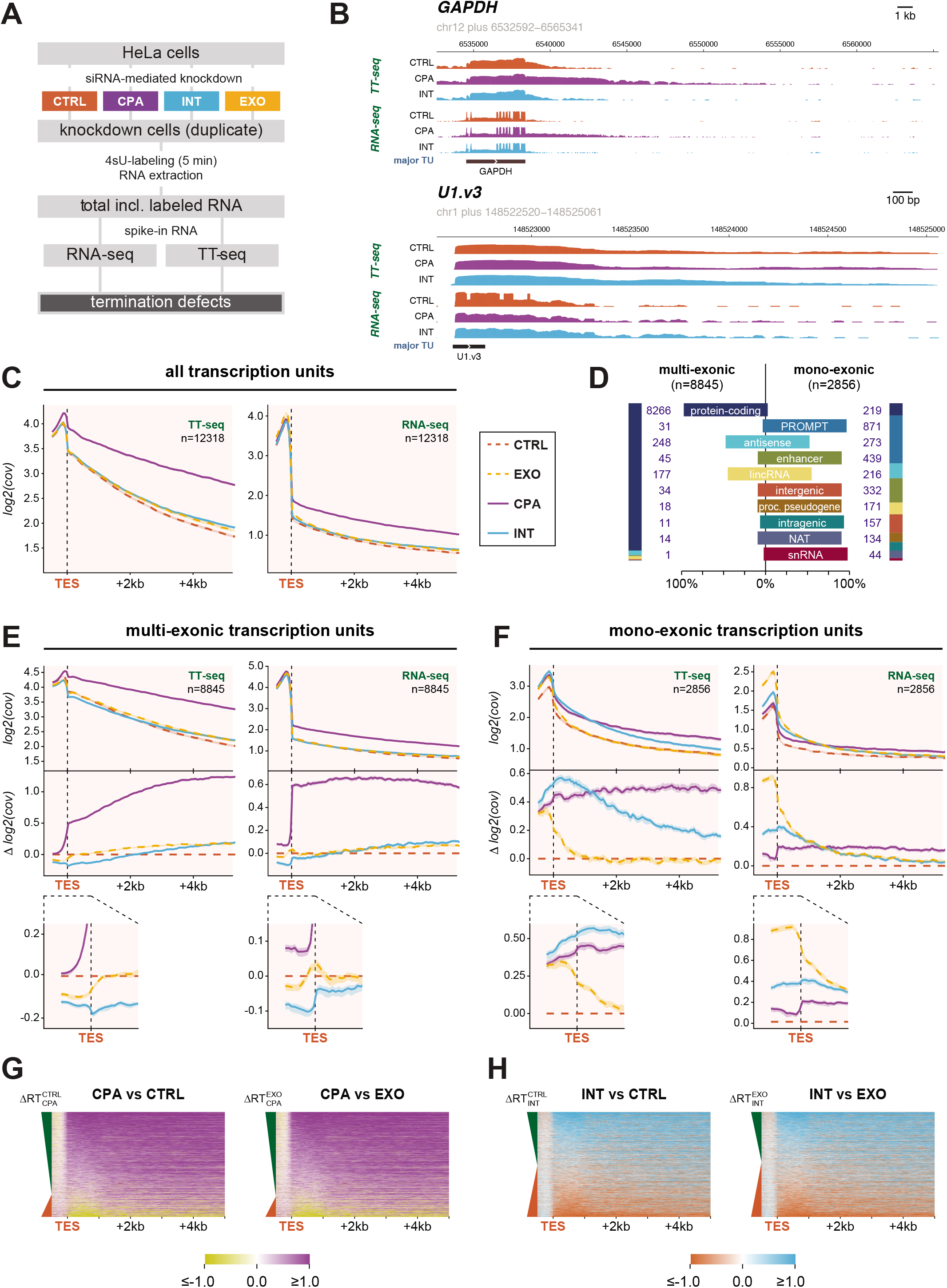
CPA and INT depletions display common and diverse global transcription termination phenotypes at TU ends. (A) Flow diagram illustrating the experimental setup for determining termination defects associated with depletion of CPSF3 and IntS11 of the CPA and INT complexes, respectively, and the control samples CTRL and EXO. (B) Genome browser views of *GAPDH* (protein-coding) and *U1.v3* (snRNA) genes and their downstream regions. Mean (of two replicates) TT-seq and RNA-seq coverage on the plus strand from control (CTRL, *red*), CPA (*purple*) and INT (*cyan*) depleted HeLa cells are shown. (C) Aggregate plots of TT-seq (*left*) and RNA-seq (*right*) data aligned to the primary transcript end site (TES) of all TUs. Mean values and 90% confidence intervals of log2-transformed coverage for 50-bp bins over all included TUs are displayed. The number of aggregated loci (n) is indicated. (D) Bar chart showing the fractions of multi- and mono-exonic TUs (in %) for the main biotypes (*middle*). The bars are annotated with the number of TUs in each set. The color code in the side bars is as in the central bar chart and represents the proportions of the respective biotypes in the multi- (*left*) and mono-exonic (*right*) TUs. (E) -(F) Aggregate plots of TT-seq (*left*) and RNA-seq (*right*) data aligned to the TES of (E) multi- and (F) mono-exonic TUs. Mean values and 90% confidence intervals of the log2-transformed Δ coverage of each of the samples vs. CTRL for the same region is shown below with enlarged plots of the region ±0.5 kb of the TES. Representation as in (C). (G) Heatmaps illustrating TES-aligned log2-transformed fold-change in mean coverage (of two replicates) in CPA depletion vs. CTRL (*left*) or vs. EXO (*right*) depletion for TUs sorted by log2-transformed difference in readthrough levels ΔRT from high to low. The green and red indicators represent the decreasing positive and negative ΔRT values, respectively. Purple indicates increased coverage and yellow indicates decreased coverage. (H) As (G) but for INT depletion vs. CTRL (*left*) or vs. EXO (*right*) depletion, respectively. Cyan indicates increased coverage and orange indicates decreased coverage. See Figure S2 for additional data related to this figure.

We next sought to assess the impact of the CPA and INT complexes on transcription termination genome-wide. Aggregate RNA coverage plots of all eligible TUs (as described in *Materials and Methods* and indicated in Figure S1E-F; FILTER +) demonstrated a bulk termination defect and associated transcriptional readthrough phenotype upon depletion of CPA, which was visible in both TT- and RNA-seq data (Figure 2C). At this analytic level, INT depletion yielded no observable general effect. However, increased signal downstream of transcript end sites (TESs) could clearly be detected upon INT depletion when analyzing TUs belonging to the PROMPT, enhancer, intragenic, intergenic and NAT categories (Figure S2B, D, F, H, I). We therefore advanced our bulk visualization by stratifying the investigated TUs into two general groups based on the absence or presence of introns in the derived transcripts, which turned out to be a defining feature for the biological responses investigated (see below). The multi-exonic category contained primarily protein-coding genes and approximately half of the antisense and lincRNA TUs, whereas the mono-exonic category consisted of the majority of PROMPT, enhancer, intergenic, intragenic and snRNA TUs as well as the remaining antisense and lncRNA TUs (Figure 2D).

As evidenced by TES-anchored aggregate plots (Figure 2E-F), multi-exonic TUs appeared suitable surrogates for protein-coding genes, whereas mono-exonic TUs generally recapitulated PROMPT and enhancer TU features (compare Figure 2E-F to Figure S2A-J). This basic division of TUs reiterated that CPA is a termination factor, acting at the TESs of both multi- and mono-exonic loci (Figure 2E-F). Recapitulating its impact on PROMPT, enhancer, intragenic, intergenic and NAT TUs, depletion of INT led to a bulk termination defect at mono-exonic TU TESs (Figure 2F). Although signal, relative to the CTRL, was also increased upstream of mono-exonic TU TESs (which was true for both CPA and INT depletion), comparison to the signal from EXO-depleted cells, which were expected to only display post-transcriptional effects, suggested clear readthrough phenotypes upon both CPA and INT depletions (Figure 2F, left panel, see bottom zoomed-in plot). However, the different shapes of the curves downstream of the TES suggested that inhibition of CPA and INT elicits distinct effects on transcription termination (discussed further below). Finally, and in clear contrast to CPA depletion, multi-exonic TUs displayed a conspicuous lowered signal immediately downstream the TES upon INT depletion (Figure 2F and S2A, discussed further below). In accordance with these conclusions based on aggregate signals, we found that of the investigated 12296 TUs, 9665 and 6289 displayed a positive TES readthrough score (ΔRT, *see Materials and Methods*) upon CPA and INT depletion, respectively (Figure 2G-H). Strikingly, however, nearly half (6133) of all analyzed TUs exhibited a negative readthrough score upon INT depletion (Figure S2G-H).

Based on these initial analyses, we conclude that CPA is a global transcription termination factor, which is not restricted to protein-coding genes. INT also appears to impact transcription termination globally. However, the effects of its depletion near TESs were not easily interpretable, calling for additional examination.

### CPA and INT depletions cause increased promoter-proximal transcription globally

The puzzling effects upstream of TESs upon depletion of both INT and CPA (Figure 2F) directed our attention to TU bodies and transcript start sites (TSSs). As expected, and due to the production of short exosome-sensitive transcripts, aggregate TT- and RNA-seq coverage plots, anchored at the major TSSs, demonstrated increased post-transcriptional signal near the promoter upon depletion of EXO for both multi- and mono-exonic TUs (Almada et al., 2013; Andersson et al., 2014a; Flynn et al., 2011; Ntini et al., 2013; Preker et al., 2008) (Figure 3A-B). In addition, these profiles revealed two prominent features: (1) depletion of both CPA and INT caused a globally increased promoter-proximal TT-seq signal for multi-as well as mono-exonic TUs, albeit more robustly for the latter category (Figure 3A-B, left panels and Figure 3C-D), and (2) for multi-exonic TUs, the aggregate TT-seq signal upon depletion of INT was indeed higher than CTRL near the promoter, but then decreased below CTRL from ~1.3 kb and downwards (Figure 3A, left bottom panel), indicating premature RNAPII termination (see below). Such depletion effects were also apparent when analyzing the underlying biotypes individually (Figure S3A-J).

**Figure 3.**
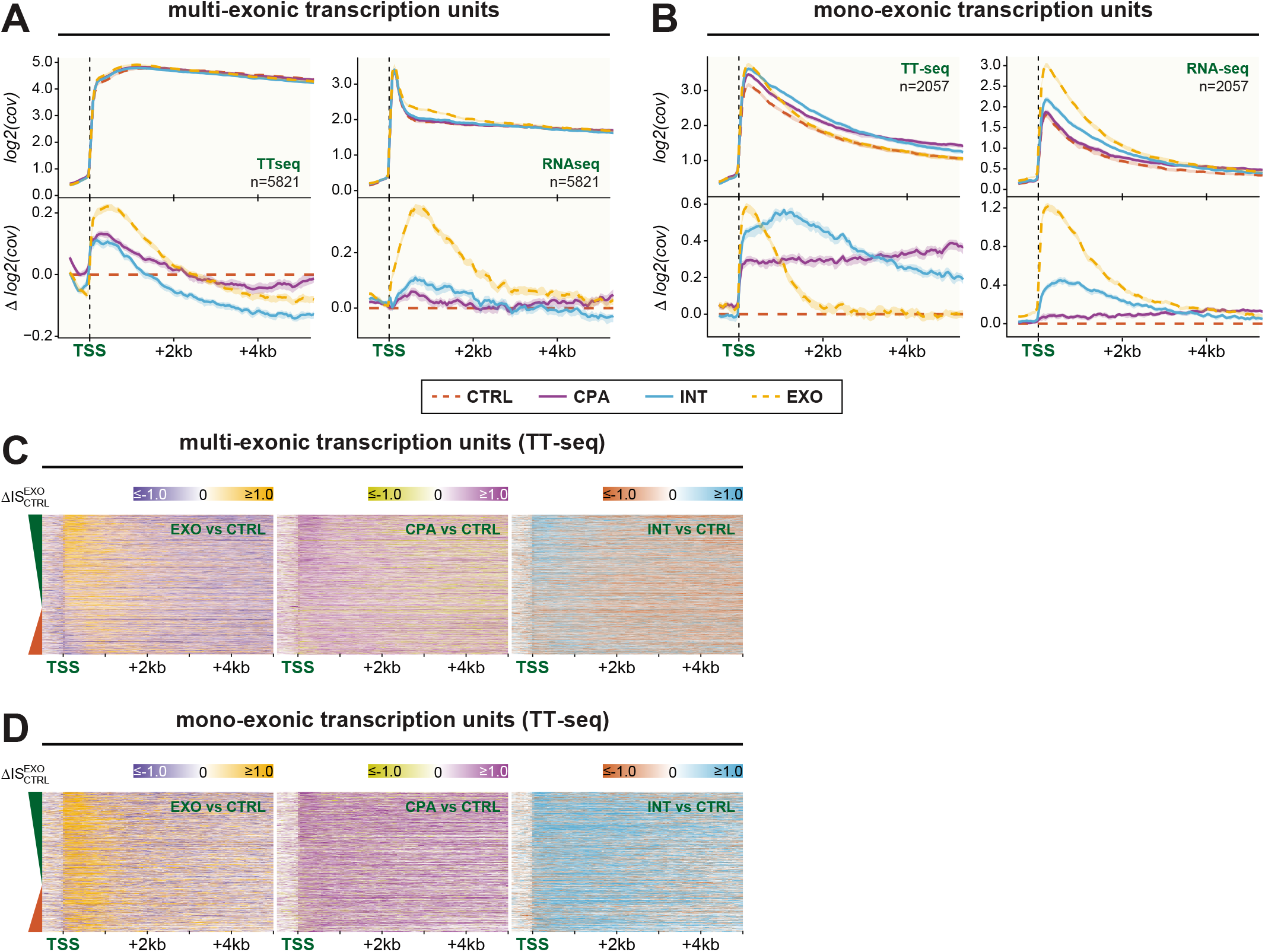
CPA and INT depletions cause increased promoter-proximal transcription globally. (A)-(B) Aggregate plots of TT-seq (*left*) and RNA-seq (*right*) data aligned to the major transcript start site (TSS) of (A) multi- and (B) mono-exonic TUs. Representation as in Figure 2E. (C)-(D) Heatmaps illustrating TSS-aligned log2-transformed fold-change in mean coverage (of two replicates) for EXO, CPA and INT depletion versus CTRL for (C) multi- and (D) mono-exonic TUs sorted by ΔIS between EXOSC3- and CTRL-depleted samples (log2-transformed difference in ‘initiation signal’ measured over the first 500 bp of the TUs) from high to low. The green and red indicators represent the decreasing positive and negative ΔIS values, respectively. Orange, purple and cyan indicate increased coverage for EXO, CPA or INT vs. CTRL, respectively. Blue, olive green and red indicate decreased coverage for EXO, CPA and INT vs. CTRL, respectively. See Figure S3 and S4 for additional data related to this figure.

Notably, the mentioned EXO and INT depletion phenotypes were also observable in the RNA-seq data, which was less pronounced for CPA depletion (Figure 3A-B, compare left and right panels). This apparently transient increase in promoter-proximal signal upon CPA depletion was confirmed by published mNET-seq data and RNA-seq data based on RNA purified from the chromatin fraction (Chr-seq) (Figure S4A-C) (Nojima et al., 2015; Schlackow et al., 2017). We note that CPA contributes to termination events near TSSs (Almada et al., 2013; Ntini et al., 2013) and that promoter-proximal pA site usage has been shown to reduce transcription initiation (Andersen et al., 2012), which together may explain the observed increase in TT-seq signal upon CPA depletion.

A global effect of INT near RNAPII promoters had not previously been recognized and we therefore aimed to validate our observations by analyzing published global run-on (GRO)-seq data from HeLa cells subjected to depletion of IntS11 (Gardini et al., 2014). Evidently, both multi- and mono-exonic TUs displayed increased promoter-proximal GRO-seq signals upon INT depletion (Figure S4D), supporting a general function for INT in restricting transcriptional output from promoters.

We conclude that both INT and CPA globally limit promoter-proximal transcription, albeit in different ways. Additionally, however, the INT complex appears to elicit and prevent transcription termination at mono- and multi-exonic TUs, respectively. To explore these seemingly contradictory activities, we focused our attention on the function of the INT complex.

### The INT complex operates to attenuate transcription in ~3kb promoter-proximal regions

Having established global activity of INT near TSSs, we set out to investigate this phenomenon in further detail using higher resolution data. We therefore conducted 3’-end-sequencing (3’-end-seq) of purified 4-thio-uridine (4sU) labeled RNA produced for 10 min in HeLa cells treated either with control siRNA (CTRL) or siRNAs directed against IntS11 (INT), EXOSC3 (EXO) or both (INT/EXO) (Figure 4A, S5A). To allow detection of both pA^+^ and pA^-^ RNA species, a fraction of each sample was treated with recombinant *E. coli* pA polymerase (ePAP) before sequencing (Wu et al., 2020) (Figure 4A). Due to size restrictions, sequenced RNA fragments were predominantly >75 nts, thus underrepresenting 3’-ends residing in promoter-proximally stalled RNAPII and hereby leaving the sequencing ‘real estate’ to other 3’-ends.

**Figure 4.**
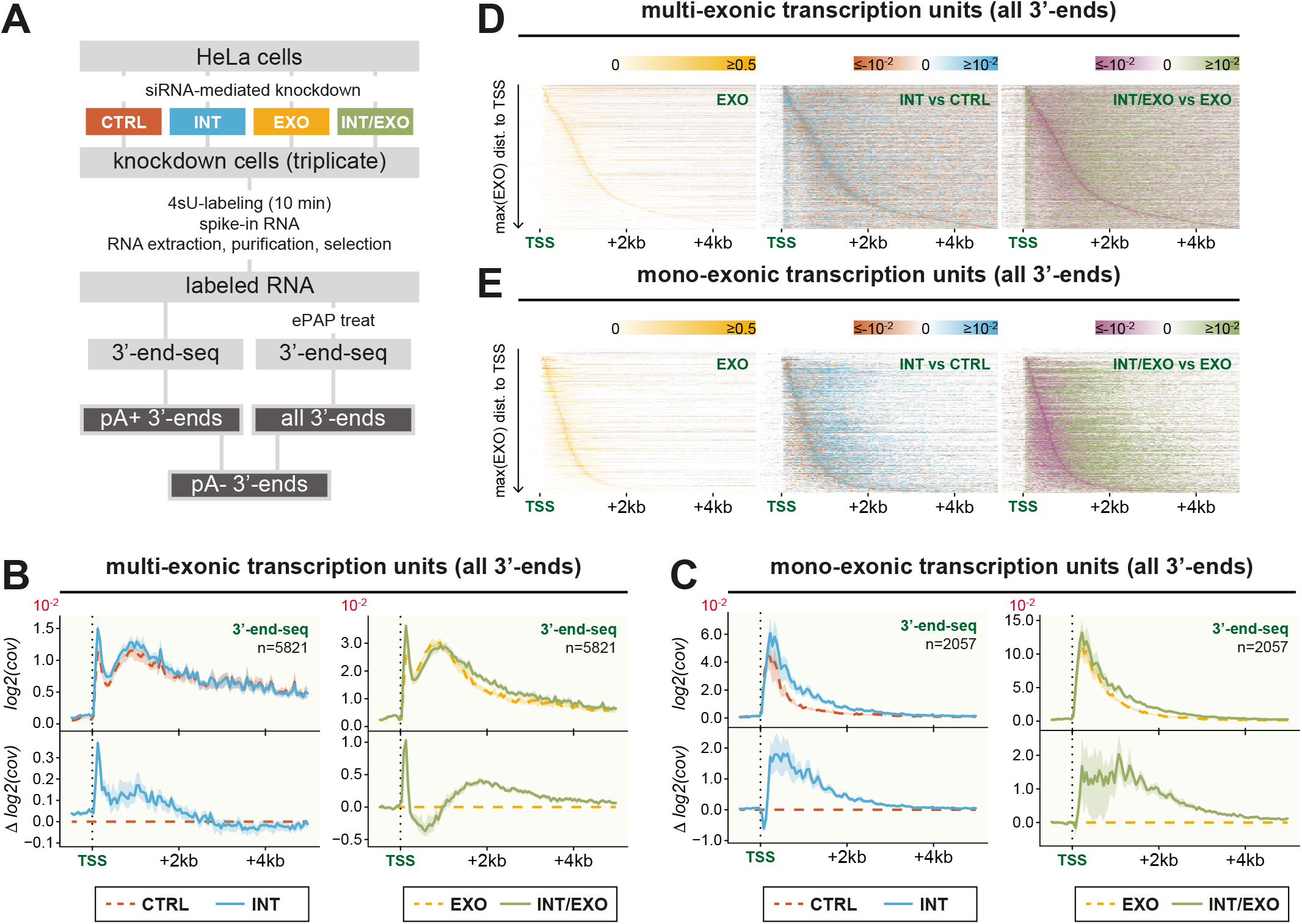
The INT complex operates to attenuate transcription in ~3kb promoter-proximal regions. (A) Flow diagram illustrating the experimental setup for determining pA^+^, all and deduced pA^-^ 3’-ends associated with depletion of INT in the background of either CTRL or EXO depletion. (B) Aggregate plots of all 3’-end-seq data (pA^+^ and pA^-^) aligned to the major TSSs of multi-exonic TUs (*top*) and log2-transformed Δ coverage for INT or EXO/INT depletions vs. CTRL or EXO depletions, respectively (*bottom*). Representation as in Figure 2E. Left panel shows coverage for CTRL (*red*) and INT depletion (*cyan*), right panel shows EXO (*orange*) and combined INT/EXO (*olive green*) depletion. (C) As (B), but for mono-exonic TUs. (D) Heatmaps of log2-transformed all 3’-end-seq data mean coverage (of three replicates) aligned at the TSS for EXO (*left*), and log2-transformed Δ mean coverage (of three replicates) for INT (*middle*) and INT/EXO (*right*) at multi-exonic TUs sorted by the distance of the maximum signal in EXO depletion to the TSS (*orange; left*). Cyan and green indicate increased signal and red and purple indicated decreased signal for the INT and INT/EXO depletions, respectively. (E) As (D), but for mono-exonic TUs. See Figure S5 for additional data related to this figure.

Gratifyingly, despite different profiles for multi- and mono-exonic TUs (see below), the 3’-end-seq data generally confirmed our previous conclusions based on TT- and RNA-seq in that there were increases in promoter-proximal and downstream signals for both multi- and mono-exonic TUs upon depletion of INT (Figure 4B-C), the general nature of which was substantiated by heatmap representations of the data (Figure 4D-E). Moreover, there was on average 3-4 fold more 3’-ends near the TSSs of mono-than multi-exonic TUs (Figure 4B-C, note scales on y-axes), despite generally lower TT- and RNA-seq coverage in mono-vs multiexonic TUs (Figure 3A-B). We suggest that this reflects generally increased promoter-proximal transcriptional termination on mono-exonic TUs, corroborated also by the declining TT- and RNA-seq levels at the same position in mono-vs multi-exonic TUs (Figure 3A-B).

We then aimed to pinpoint more precisely where INT operates. Since INT action is expected to produce non-adenylated 3’-ends (Guiro and Murphy, 2017), we subtracted pA^+^ from total 3’-end signals to estimate pA 3’-end quantities (Figure S5B-C), which turned out to be the dominating species (compare Figure 4B-C to S5B-C and 4D-E to S5D-E). The aggregate TSS-anchored pA 3’-end-seq profiles were notably different between multi- and mono-exonic TUs (compare Figure S5B to S5C), reflecting the diverse length and intron-content of these two TU categories. Mono-exonic TUs displayed a single TSS-proximal signal peak (Figure 4C, S5C, left panels), whereas multi-exonic TUs displayed two peaks (Figure 4B, S5B) of which the first could be linked to TSS positioning (data not shown) and likely reflected a ‘shoulder’ of stalled RNAPII (Core and Adelman, 2019; Jonkers and Lis, 2015), while the second was a consequence of the presence of introns (data not shown). Despite these different profiles, the 3’-end signal increased and extended further away from the TSS upon depletion of INT for both mono- and multi-exonic TUs (Figure 4B-C, S5B-C, left panels), which again was underscored by heatmap representation of the data (Figure 4D-E, S5D-E). Moreover, EXO depletion, expected to stabilize INT-generated 3’-ends, generally led to a 2-3 fold increase in 3’-end-seq signals (Figure 4B-C, S5B-C, compare left to right panels). Somewhat surprisingly, co-depletion of INT and EXO shifted those 3’-ends a few kb downstream compared to the EXO depleted sample (Figure 4B-E, S5B-E), such that the curves of the aggregate plots merged at app. 4 kb downstream of the TSS for both mono- and multi-exonic TUs (Figure 4B-C, S5B-C, right panels). Thus, although INT has a prominent and genome-wide activity in generating 3’-ends within 3 kb from promoters, alternative RNA 3’-ends are still produced further downstream in its absence.

Based on the combined impression from TT-, RNA-, GRO- and 3’-end-seq, we conclude that INT operates within a ~3 kb zone near most, if not all, RNAPII promoters and with its activity being most prominent within the first 1 kb. Notably, although early termination is inhibited by depletion of INT, the ensuing decline of promoter-distal signals indicates that RNAPII transcription still ceases at downstream positions (≤4 kb from promoters) indicative of yet additional termination mechanisms.

### INT depletion exposes non-productive RNAPII transcription events

To address whether an increased usage of premature pA sites might contribute to downstream transcription termination in the absence of INT, we plotted the distribution of pA^+^ RNA 3’-ends near TSSs of multi- and mono-exonic TUs (Figure 5A-B). Depending on the exact distance to the TSS, these 3’-ends constituted from one-tenth to one-third of the total 3’-end signal in the control and INT depleted samples (compare left panels in Figure 4B and 4C with those of 5A and 5B), which increased upon EXO co-depletion consistent with the production of unstable RNA (Chiu et al., 2018; Ogami et al., 2017; Wu et al., 2020) (compare right panels in Figure 4B and 4C with those of 5A and 5B). Overall, the bulk pA^+^ signal appeared more distal to the TSS compared to that of the total RNA 3’-end signal (compare Figure 5A-B to 4B-C, left panels), reflecting pA site-dependent transcription termination events. Indeed, alignment to the first putative pA signal within the promoter-proximal 5 kb (based on presence of AWUAAA) and which did not coincide with the major TES, revealed a substantial amount of polyadenylated species deriving from these sites (Figure 5C-D). These pA^+^ 3’-end signals increased 4-to 5-fold in EXO samples (Figure 5C-D, compare scales on right and left panels), illustrating their generally unstable nature. Importantly, such pA site usage was robustly increased upon INT depletion (Figure 5C-D, left panels), strongly suggesting that a share of the RNAPII complexes evading termination by INT instead fall prey to pA site-dependent termination. In the case of multi-exonic TUs, this increased usage of premature pA sites may well explain the observed drop below the CTRL signal in INT depleted TT-seq samples (Figure 3A), which was clearly visible downstream of the putative pA sites (Figure 5E, left panel). We take these observations to suggest that an important promoter-proximal activity of the INT complex is to terminate a fraction of early elongating polymerases, which would otherwise fail to engage in productive elongation and terminate at alternative positions such as cryptic pA sites. Such termination might be part of an RNAPII checkpoint where promoter-proximal stalling contributes time to configure RNAPII for efficient transcription (see *Discussion*).

**Figure 5.**
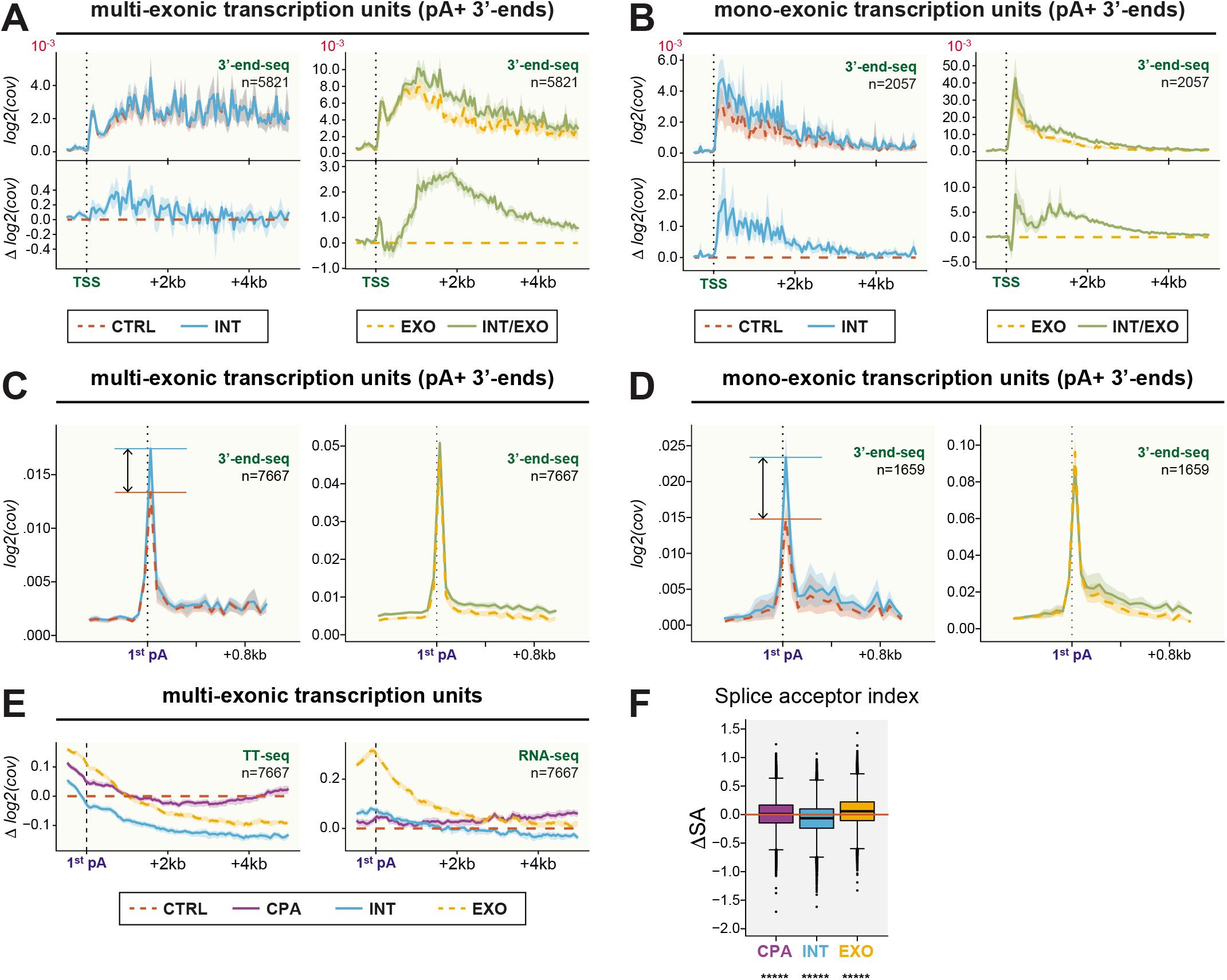
INT depletion exposes non-productive RNAPII transcription events. (A) Aggregate plots as in Figure 4A, but of pA^+^ 3’-end-seq data. (B) As (A), but for mono-exonic TUs. (C) Aggregate plots of pA+ 3’-end-seq data aligned at the first encountered putative pA signal (1^st^ pA) of multi-exonic TUs. Representations as in (A). Horizontal lines in corresponding colors indicate the peak height and the arrows indicate the difference in peak height. (D) As (C), but for mono-exonic TUs. (E) Aggregate plots of the log2-transformed TT-seq (*left*) and RNA-seq (*right*) Δ coverage of CPA (*purple*), INT (*cyan*) and EXO (*orange*) depletions vs. CTRL aligned as in (C). Representations as in (C). Equivalent plots for mono-exonic TUs are shown in Figure S5D. (F) Box plot displaying the changes in splice acceptor ratio (ΔSA) for CPA (*purple*), INT (*cyan*) and EXO (*orange*) depletions relative to CTRL for all multi-exonic TUs. Box limits represent the first and third quartiles, the band inside the box is the median. The ends of the whiskers extend the box by 1.5 times the interquartile range. The means of all three ΔSA distributions were significantly different from zero (Wilcoxon test, *****p < 2.2e-16). See Figure S5 for additional data related to this figure.

If the INT complex functions at an early RNAPII checkpoint, co-transcriptional RNAPII-dependent RNA processing, such as pre-mRNA splicing, might also be affected upon INT depletion. To address this possibility, we assessed splicing competence by using the RNA-seq data to calculate a ‘splicing ratio’ based on the number of spliced vs. unspliced reads mapping to all detected splice acceptors (splice donors were not included because the measure was skewed for TSS-proximal splice donors due to increased promoter-proximal transcription). When comparing the distributions of these scores relative to the CTRL, a net decrease in splice acceptor usage upon INT depletion became evident (Figure 5F). The same was true for splice donors of internal introns (data not shown). Due to the long half-lives of snRNPs (Baillat et al., 2005; Yamamoto et al., 2014), we considered it unlikely that this effect would be caused by altered snRNA levels under INT depletion conditions. This notion was supported by our finding that sampled sets of multi-exonic TUs, which either only displayed (1) a minor variation in, (2) increased or (3) decreased splice site usage upon INT depletion (Figure S5G, I, K) still exhibited TT-seq profiles and premature pA site usage similar to that of all multi-exonic TUs (compare Figure S5H, J, L to 5C).

Taken together, we suggest that INT depletion exposes non-optimal, possibly ill-configured, RNAPII complexes to continued transcription, which results in inefficient co-transcriptional RNA processing and transcriptional elongation.

### INT activity correlates inversely with transcriptional output

The effects of INT depletion on promoter-proximal TT-, RNA- and 3’-end-seq signals were more prominent for mono-relative to multi-exonic TUs (Figures 3A-D, 4B-E). Since mono-exonic TUs are generally expressed at lower levels than multi-exonic TUs (Figure S1G), we wondered whether INT preferentially dampens expression from lowly expressed TUs (weak promoters). To address this question, we divided all eligible (i.e. filtered for incoming upstream signal) TUs into six groups based on their mean expression levels as measured by TT-seq coverage under control conditions, and, as expected, found the top and bottom groups dominated by multi- and mono-exonic TUs, respectively (Figure 6A). We then plotted the aggregate differences in TT-seq signals between factor depleted and CTRL samples anchored to TU TSSs (Figure 6B), which revealed a strong negative correlation between INT depletion effects and TU expression levels (Figure 6B, left panel). This relationship was general for both multi- and mono-exonic TUs (Figure S6A-B), and a similar correlation was expectedly seen for EXO depletion samples (Figure 6B, right panel) (Lloret-Llinares et al., 2018). Moreover, both profiles were also visible when plotting our RNA-seq data (Figure S6C, left and right panels). Taken together with the 3’-end-seq data comparing EXO depleted samples with or without co-depletion of INT (Figure 4A-B, right panels), this data is in accordance with a large share of promoter-proximal exosome sensitive RNAs being INT terminated. A similar tendency was observable in TT-seq data upon depletion of CPA, but not as sharply defined (Figure 6B, middle panel) and lost in the RNA-seq data (Figure S6C, middle panel).

**Figure 6.**
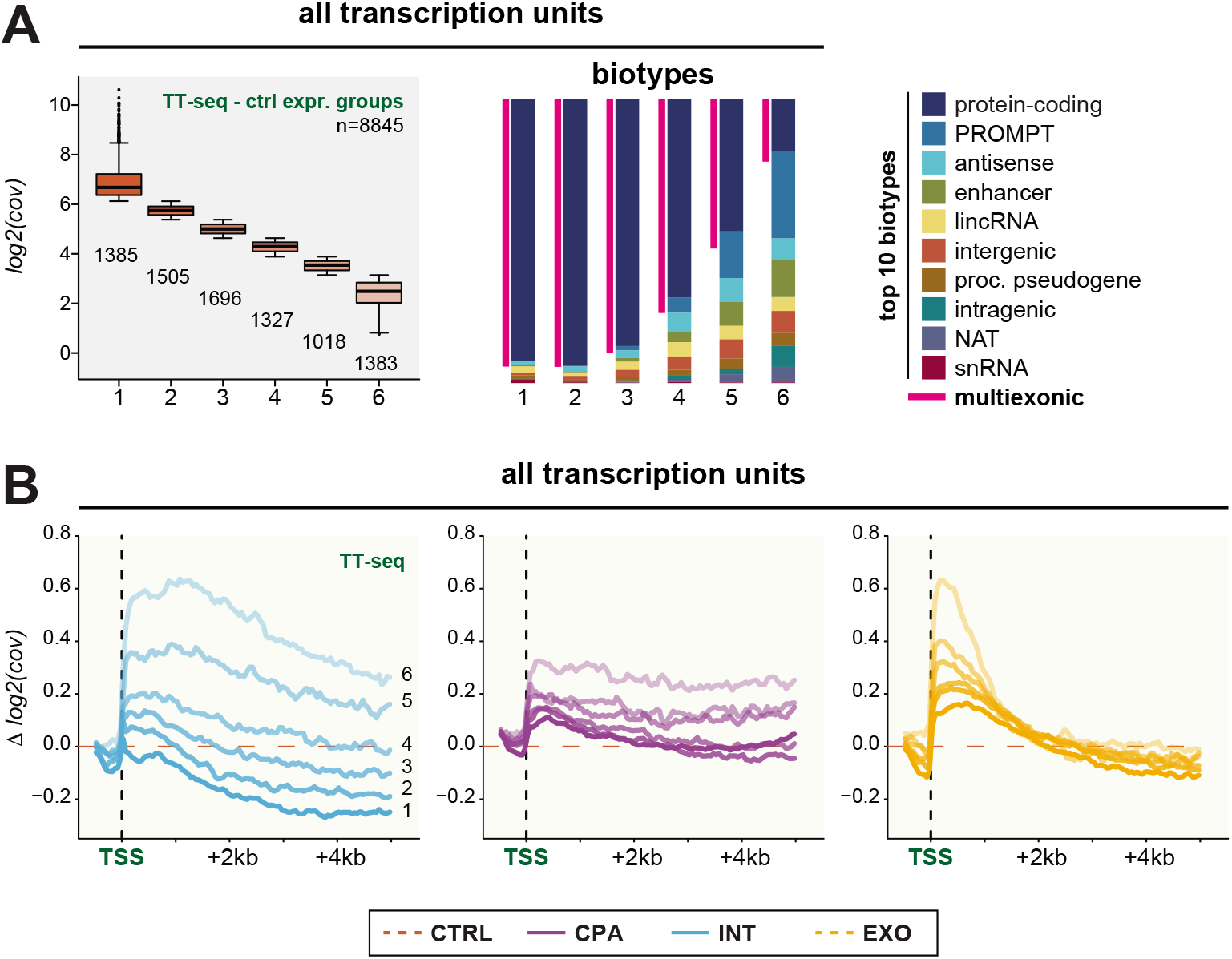
INT activity correlates inversely with transcriptional output. (A) (*Left*) Box plot displaying expression levels (measured as log2-transformed mean TT-seq coverage over the locus) for six expression-based TU groups (1-6) based on expression in CTRL samples. Box limits represent the first and third quartiles, the band inside the box is the median. The ends of the whiskers extend the box by 1.5 times the interquartile range. Outliers are show in black. The number of TUs in each group is indicated. (*Right*) Bar charts showing composition fractions of the major biotypes for the gene groups 1-6, respectively. The magenta bars indicate the proportion of multi-exonic TUs for each gene group. (B) Aggregate plots of the log2-transformed TT-seq Δ coverage of INT (*cyan, left*), CPA (*purple, middle*) and EXO (*orange, right*) depletions vs. CTRL aligned at the major TSS. Mean values of log2-transformed Δ coverage for 50-bp bins are displayed. The line color intensity (high to low) corresponds to the gene group 1 to 6, respectively. See Figure S6 for additional data related to this figure.

We conclude that the activity of the INT complex is relatively more prominent on lowly expressed TUs, which in turn results in rapid exosomal turnover of the expressed transcripts. This may serve to considerably suppress ‘background’ transcription from pervasively transcribed promoters.

### snRNA TUs exploit INT-sensitive transcription for their stable RNA production

Having established a role for the INT complex as a generic terminator of promoter-proximal transcription begged the question how this can be reconciled with INT function in the 3’-end processing of snRNAs? Comparing the TES-anchored profiles of TT-, RNA- and 3’-end-seq data from snRNA TUs with general mono-exonic TUs, such as those expressing labile PROMPTs and eRNAs, did at first glance not reveal any major differences (Figure S2B, S2D, S2J, 7A-B, S7A-B). In both cases, INT depletion led to transcriptional readthrough and the resulting ‘3’-extended RNAs’ were sensitive to exosome depletion. However, closer inspection revealed distinct, and INT-sensitive, 3’-ends from within snRNA genes (Figure S7C-F), whereas other mono-exonic TUs generally displayed 3’-ends scattered over a wider area (Figure S7G-H). To understand this in greater detail, we mapped the genomic positions of prominent INT-sensitive 3’-ends – i.e. high-signal 3’-ends that were significantly ‘downregulated’ upon depletion of INT (see *Materials and Methods* for details) and found snRNA TUs to be strongly enriched (Figure 7C, S7I). Additionally, we found a number of INT-sensitive 3’-ends falling within multi-exonic and non-snRNA mono-exonic TUs categories (Figure 7D, S7J). In the case of multi-exonic TUs, these INT-sensitive sites coincided with TESs (Figure 7E, S7A, S7K), which we interpret as an indirect effect of the observed lowered RNAPII processivity over these TUs upon INT depletion thus leading to fewer polyadenylation events (e.g. Figure 2E and S7A).

**Figure 7.**
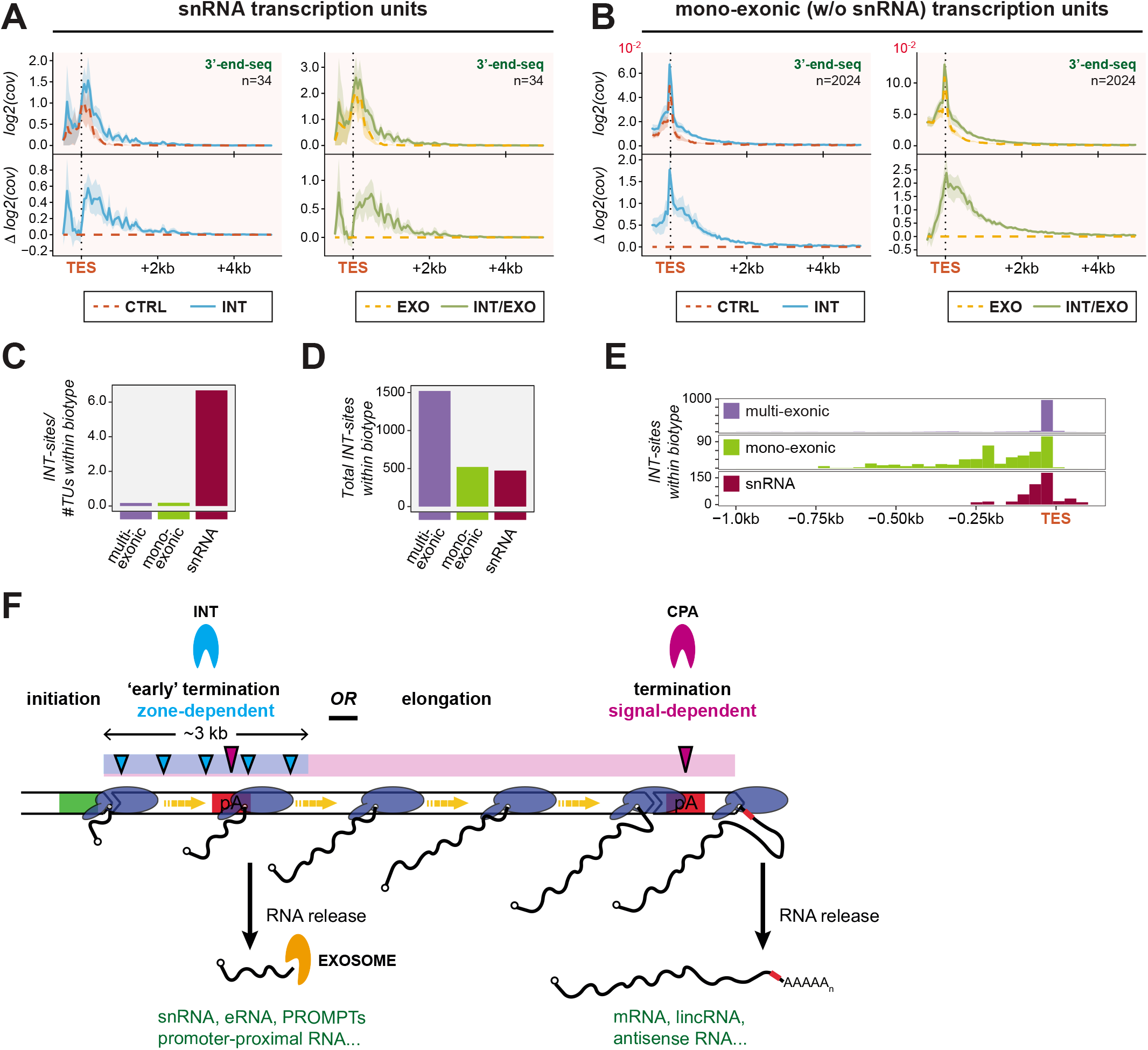
snRNA TUs exploit INT-sensitive transcription for their stable RNA production. (A) Aggregate plots as in Figure 4A, but of all 3’-end-seq data anchored to the TES of snRNA TUs. (B) As (A), but for mono-exonic TUs excluding snRNAs. (C) Relative distribution of INT-sensitive 3’-end sites (INT-sites) falling within multi-exonic, mono-exonic (excluding snRNA) and snRNA TUs. See Figure S7I for stratification into the underlying biotypes. (D) Distribution of INT-sensitive sites falling within multi-exonic, mono-exonic (excluding snRNA) and snRNA TUs. See Figure S7J for stratification into the underlying biotypes. (E) Distribution of INT-sensitive sites relative to the TES in multi-exonic (*top*), mono-exonic (excluding snRNA; *middle*) and snRNA (*bottom*) TUs. See Figure S7K for stratification into the underlying biotypes. (F) Model illustrating the proposed functional spaces for the INT and CPA complexes. See text for details. See Figure S7 for additional data related to this figure.

Manual alignment of INT-sites in snRNA TUs indicated motifs resembling the 3’-box downstream of the mapped INT-sensitive cleavage sites, which reside a few nucleotides downstream of the mature snRNA 3’-ends (Figure S7L). Due to the highly degenerate nature of these putative 3’-boxes, it was not possible to conduct a sequence-based search for these elements outside of snRNA genes. However, the identification of additional INT-sensitive 3’-ends within non-snRNA mono-exonic TUs revealed a few other loci, some expressing relatively stable RNAs, with a similar motif (Figure S7M).

Based on these observations, we propose that INT generally stimulates RNAPII termination by cleaving the nascent RNA in a largely sequence-independent manner within a promoter-proximal zone, which produces 3’-ends readily accessible to the RNA exosome. However, INT may be specifically engaged to conduct site-specific cleavages directed by 3’-box-like sequence elements. This, in turn may allow the produced RNA to deflect the RNA exosome and become a stable species (Coy et al., 2013), *see Discussion*).

## DISCUSSION

Traditionally, transcription termination has been considered to be the process ‘finalizing’ full-length gene transcription and recycling RNAP for a new round of initiation. It was suggested that RNAPII termination requires cleavage of the nascent transcript, relying on the CPA and INT complexes to act at the ends of the major protein-coding and snRNA gene classes, respectively. Although this is still correct, recent findings that transcription termination occurs pervasively both within and outside of conventional genic regions have spurred a renewed interest in how such promiscuous termination is triggered and which function(s) it may possibly exert (Core and Adelman, 2019; Kamieniarz-Gdula and Proudfoot, 2019; Porrua and Libri, 2015; Proudfoot, 2016). Investigation of these phenomena requires factor perturbation and unbiased in-depth bioinformatic analysis of different types of genome-wide data. Utilizing such RNA biotype-agnostic approaches, our results demonstrate that both CPA and INT operate globally over the human genome to terminate conventional as well as pervasive transcription. In doing so, rather than exhibiting gene class-specific activities, these complexes react to general features of the DNA/RNA template: CPA activity is signal-dependent and can be accounted for by the usage of pA sites, while INT activity is generally sequence-independent and occurs in TSS-restricted regions of ~3kb (Figure 7F). Collectively, CPA and INT therefore account for a large share of RNAPII transcription termination in the human genome. Moreover, the possible cooperation of CPA and INT, within the same DNA elements, suggests a parsimonious solution to previous conflicts over the exact termination pathways of human lncRNA transcription (Almada et al., 2013; Andersson et al., 2014b; Lai et al., 2015; Ntini et al., 2013; Schlackow et al., 2017; Wu et al., 2020). Finally, our analyses propose an unprecedented role of early termination to the quality control of RNAPII by preventing its non-productive transcription of both mono- and multi-exonic TUs.

Whereas a global activity of CPA in transcription termination had been reported (Almada et al., 2013; Ntini et al., 2013), the general genome-wide activity of INT, observed here in many regions that are devoid of canonical 3’-box sequence motifs, was unexpected. Previous (Hernandez 1985, Baillat and Wagner 2015, Guiro and Murphy 2017 and recent (Elrod et al., 2019; Tatomer et al., 2019) reports suggested that INT assists in the termination of transcription downstream of snRNA genes as well as at a subset of promoter-proximally stalled RNAPII. However, our data demonstrate that these reported cases represent only a small fraction of genomic regions exposed to INT activity. Instead, we suggest that transcription termination by INT occurs promiscuously in ~3 kb broad promoter-proximal regions and that snRNAs, and stably stalled RNAPII, present exceptionally strong substrates for INT cleavage. INT activity is visible for both mono- and multi-exonic TUs, although less so for the latter category, likely due to the presence of overlapping full-length transcription events. Moreover, promoter-proximal termination by INT is coupled to RNA degradation by the exosome for both TU categories.

It is currently unclear how the observed general INT activity is restricted to the promoter-proximal 3 kb region. Available occupancy data obtained by ChIP-seq indeed demonstrate an accumulation of INT in the 5’-region of genes (Gardini et al., 2014; Stadelmayer et al., 2014). This relates to the question of how INT is recruited to elongating RNAPII and how its cleavage activity is regulated. Multiple RNAPII-associated factors are exchanged within the promoter-proximal region, and some of these may facilitate INT binding, whereas others can compete with INT for RNAPII interaction. The outcome of such competition between positive (elongation-stimulatory) and negative (pausing- and termination-stimulatory) factors for RNAPII association likely decides the fraction of terminated complexes at the beginning of TUs. It was reported that INT co-immunoprecipitates with the negative elongation factor (NELF) (Yamamoto et al., 2014), and that INT itself recruits the positive super elongation complex (Gardini et al., 2014). Although it requires further investigation to understand this complex interplay of factors, we suggest that INT targeting might generally depend on RNAPII velocity, which is low at the beginning of genes (Jonkers et al., 2014; Saponaro et al., 2014), the availability of nascent RNA that is not yet packaged with other proteins, and on the phosphorylation status of RNAPII and its associated factors.

Regardless the details of INT recruitment and activation, we suggest that the defined INT sensitive promoter-proximal zone provides for quality control of non-productive RNAPII transcription. This is illustrated by our data in two ways. In one line of inquiry, we report evidence that early INT-dependent transcription termination is relatively more efficient when TU expression is low (Figure 6, S6). Consistently, transcripts deriving from weak promoters were also reported to be relatively more exosome-sensitive (Lloret-Llinares et al., 2018). Therefore, such transcription termination-coupled RNA decay may generally serve to suppress the substantial fraction of genomic transcription, which does not exceed a certain expression threshold and consequently is less likely to be functionally meaningful. Increased transcription overcomes such general dampening, possibly by exhausting available INT components, thereby improving the yield of stable transcripts.

The importance of INT-dependent termination of non-productive transcription can also be inferred from the apparent RNAPII processivity defects that we detected beyond the 3 kb promoter-proximal zone on multi-exonic genes upon INT depletion (Figure 2E, 3A). An observed processing defect at a subset of protein-coding gene TESs upon INT-inactivation was previously rationalized by the presence of 3’-box elements at the affected loci (Stadelmayer et al., 2014). However, the definition of the 3’-box motif used in that study differed from the one defined previously (Hernandez, 1985) (‘AAAAACAGACC’ vs ‘GTTTN_0-3_AAARN_2_AGA’), and we could not establish this connection in our data using either of the two motifs (data not shown). Instead, we suggest that the apparent ‘defect’ derives from the lack of promoter-proximal INT activity, leading to the unsolicited elongation of RNAPII complexes, evading an early quality control that is normally contributed by INT. This is because we observe aberrant usage of TES pA-sites in INT-depletion conditions (Figure S7A, 7E, data not shown). Additionally, premature pA-site usage is increased upon INT depletion (Figure 5C, E), which is consistent with the release of a subset of elongation ‘incompetent’ RNAPII in the absence of a prior INT-controlled checkpoint. Non-optimal RNAPII performance is also indicated by the decreased splicing of transcripts upon INT depletion (Figure 5F), suggesting that although the enzyme is still transcribing DNA its function in co-transcriptional RNA processing is affected, maybe because not all factors required for co-transcriptional splicing are loaded.

Finally, provided that promoter-proximal INT activity is an important contributor in terminating pervasive transcription, how do snRNA TUs, but not the TUs producing labile non-coding RNAs, manage to produce stable transcripts? Based on the overall exosome sensitivity of transcripts produced from within and downstream of snRNA TUs (Figure S2J, S3J, 7A), these appear to abide to the same INT-induced quality control as all other loci. However, we suggest that the presence of INT consensus binding sites (aka 3’-box elements) may provide an important difference. Indeed, there are well-defined INT-sensitive cleavage sites downstream of the mature snRNA 3’-ends (Figure 7C-E and S7C-F), which are not generally present at PROMPT, eRNA or otherwise mono-exonic TUs (Figure 7E and S7G-H), and which correlate with the presence of 3’-box elements (Figure S7I). We propose that snRNA TUs are outliers to the general scenario in that they allow promoter-proximal INT activity to become a constructive incident, leading to the production of stable RNA.

In summary, we propose a model where the INT complex serves a quality control function by terminating polymerases that fail to enter productive elongation. If such complexes are allowed to continue, as in the absence of INT, at least a fraction of them appears to be ill-configured to support splicing and avoid termination by premature pA sites. We note that RNA processing defects have also been reported upon NELF inactivation (Yamamoto et al., 2014), suggesting that this complex might cooperate with INT to provide quality control. This is consistent with the view that NELF extends the promoter-proximal duration of RNAPII, opening a time window for the assembly of an activated RNAPII elongation complex, that contains all factors required for processive transcription and mature mRNA production.

## ACKNOWLEDGMENTS

We thank Peter Refsing Andersen for critical comments to the manuscript. T.H.J. was supported by the European Research Council (EURECA advanced investigator grant agreement no. 339953), the Lundbeck Foundation and the Novo Nordisk Foundation. P.C. was supported by the Deutsche Forschungsgemeinschaft (SFB860, SPP1935, EXC 2067/1-390729940), the European Research Council (TRANSREGULON advanced investigator grant agreement no. 693023), and the Volkswagen Foundation.

## AUTHOR CONTRIBUTIONS

Søren Lykke-Andersen: data curation; principal bioinformatic analyses; data interpretation; manuscript writing.

Kristina Žumer: TTseq/RNAseq data collection; data interpretation; curation of published datasets; manuscript writing.

Ewa Molska: performed and validated factor depletions; performed experiments for TTseq/RNAseq; collected cell lysates.

Guifen Wu: performed experiments and collected data for 3pSeq.

Jérôme O. Rouvière: curation of published datasets; bioinformatic analyses.

Manfred Schmid: initial 3pSeq data curation; bioinformatic analyses.

Carina Demel: assisted with TU annotation and implementation.

Björn Schwalb: assisted with TU annotation and bioinformatic analysis.

Patrick Cramer: supervision; data interpretation; manuscript writing; funding. Torben Heick Jensen: supervision; data interpretation; manuscript writing; funding.

## MATERIALS AND METHODS

### Cell culture and RNA interference

For the TT- and RNA-seq experiments, HeLa Kyoto cells were grown in Dulbecco’s modified eagle medium (Invitrogen) supplemented with 10% fetal bovine serum and 1% penicillin/streptavidin at 37°C and 5% CO_2_. RNA interference was used for depletion of the desired protein factors (siRNA sequences are shown below). Two rounds of transfections were performed. The first transfections were performed using SilentFect (Biorad) (final dilution 1:1000) in RPMI 1640 media 24 hours after seeding the cells. Two days after the first transfection, the media was changed, and the transfection was repeated using Lipofectamine 2000 (final dilution 1:1000) in RPMI 1640 media. siRNAs were used at a final concentration of 15 nM of each siRNA. 24 hours after the second round of transfections, cells were labelled for 5 min with 4-thiouridine minutes and harvested.

3’-end-seq experiments were conducted as described (Wu et al., 2020).

The following protein were depleted (siRNA sequences in brackets):

CTRL (EGFP for TT- and RNA-seq experiments: GACGUAAACGGCCACAAGUdTdT/ ACUUGUGGCCGUUUACGUCdTdT and LUC for 3’-end-seq experiment: CUUACGCUGAGUACUUCGAdTdT / UCGAAGUACUCAGCGUAAGdTdT), CPA (CPSF3: CUUGUGGCCGUUUACGUCdTdT / CACAGUCACGACUAGGUCAdTdT), INT (IntS11: AGACAACAAGCAUGCGAAAdTdT / CAUCAAGCAUGCAGAGAAAdTdT) and EXO (EXOSC3: CACGCACAGUACUAGGUCAdTdT/UGACCUAGUACUGUGCGUGdTdT).

### Western blotting

Knockdown efficiency was confirmed by western blotting of whole-cell extracts. Pelleted cells were resuspended in RSB100 (100 mM Tris pH 7.5, 100 mM NaCl, 2.5 mM MgCl2, 0.5% Triton-X100) supplemented with proteinase inhibitor followed by sonication. Cell debris was removed by centrifugation at 4000 g and 4°C for 15 minutes. Samples were separated by 10% denaturing SDS-PAGE and transferred to PVDF membrane (Milipore). Membranes were blocked in 5% skimmed milk powder (SMP) in PBS for 30 minutes and incubated with primary antibodies (see below) in 5% SMP in PBS. The membranes were subsequently washed three times for 10 minutes in PBS with 0.05% Tween20 and incubated with horseradish peroxidase (HRP) conjugated anti-rabbit or anti-mouse secondary antibody (Dako) in 5% SMP in PBS, washed again and exposed using Supersignal West Femto Substrate (Thermo Scientific).

The following proteins were detected by western blotting (antibodies in brackets) CPSF3 (Sigma-Aldrich, HPA04657), IntS11 (Sigma-Aldrich, HPA029025), EXOSC3 (Proteintech Europe, 15062-1-AP), SRRT (Abcam, ab55822), XRN2 (Bethyl, A301-103A) and TUBULIN (Rockland, 200-301-880).

### TT- and RNA-seq library construction

TT- and RNA-seq experiments were done in duplicate. The TT-seq protocol was performed as previously described (Schwalb et al., 2016; Zylicz et al., 2019) with some modifications described here in brief. The spike-in mix, containing three 4sU-labeled and three unlabeled RNAs, was synthesized with MEGAscript T7 Transcription Kit (Thermo Fisher). For synthesis of the 4sU-labeled spike-ins 1/10 of UTP in the transcription reactions was replaced with 4sUTP (Jena BioScience).

RNA lysates in Trizol were thawed on ice and 400 pg of synthetic spike-in mix was added per 1 million cells to each of the samples. The total RNA was then purified according to the manufacturer’s protocol. 4sU-labeled RNA was isolated from 600 μg total RNA, split into two equal aliquots for the isolation. These were sheared in Covaris MicroTubes with the Covaris S220 System (Covaris) for 10 s with settings 100 W and 1% duty cycle. These were then pooled and 2 μg of sonicated total RNA was collected for total RNA fraction (RNA-seq) and stored at −80°C until proceeding with purification and NGS library preparation. The rest of the sheared RNA was biotinylated with EZ-Link HPDP-Biotin (Thermo Fisher) in two biotinylation reactions per sample (300 μg RNA in 10 mM Tris pH 7.5; 1 mM EDTA, 20 % DMF; and 200 μg/ml HPDP-biotin) that were incubated for 1.5 h at 24°C. Excess biotin was removed by extracting the biotinylated RNA with chloroform and precipitation with isopropanol. 4sU-labeled RNA was isolated by affinity purification with paramagnetic streptavidin μMACS MicroBeads (Miltenyi). Biotinylated-4sU-labeled RNA was incubated with the paramagnetic beads for 15 min at room temperature with gentle agitation. The beads were then loaded onto MACS columns (Miltenyi). The columns were washed three times with wash buffer (100 mM Tris pH 7.5; 10 mM EDTA; 100 mM NaCl; and 0.1% Tween 20) at 65°C and three times with wash buffer at room temperature. The 4sU-labeled RNA was eluted from the columns with 100 mM DTT. The eluted 4sU-labeled RNA and reserved total RNA were purified with the miRNeasy Micro kit (Qiagen) according to the manufacturer’s protocol with on-column DNase I treatment. NGS libraries were generated from 100 ng and 600 ng of isolated newly transcribed RNA and total RNA fractions, respectively, using the TruSeq Stranded Total RNA Library Prep Kit (Illumina) according to the manufacturers protocol. The sample libraries were pooled and sequenced on the HiSeq2500 (Illumina) using PE50 mode.

### TT- and RNA-seq data processing

Raw sequence reads from TT- and RNA-seq experiments were mapped to the reference genome using STAR aligner (Dobin et al., 2013) and putative PCR duplicates were removed using samtools (Li et al., 2009). The reference genome was derived from the catenation of fasta files with the human genome release GRCh38 and six spike-in RNAs (Schwalb et al., 2016). Only uniquely mapping reads were kept for downstream analyses.

Size-factors for sequencing depth normalization were obtained using the DESeq2 algorithm (Love et al., 2014) from HTSeq count data (Anders et al., 2015) on a merged GenCode v21 annotation.

The reads mapping to spike-ins were used to estimate antisense bias, and TT- and RNA-seq data were corrected accordingly (Schwalb et al., 2016).

TT-seq data were additionally corrected for contaminating unlabeled RNA based on spike-in RNA estimations (Schwalb et al., 2016). Moreover, for TUs with data-supported splicing (multi-exonic) the level of spliced RNA was estimated for each locus and subtracted to provide a conservative measure of newly transcribed RNA.

The experiments were done in duplicates in two batches and the data were batch corrected using the limma package (Ritchie et al., 2015) leading to a good coclustering of replicates in PCA analysis based on log2-transformed mean coverage over the individual loci (data not shown).

### *De novo* HeLa transcriptome annotation

We generated a transcriptome annotation based on our experimental data. In brief, CTRL and EXO depletion samples were used to identify transcribed units (TUs). The EXO depletion stabilizes unstable transcripts such as PROMPTs, eRNAs and NATs. TT-seq data were used to identify regions of continuous transcription based on a hidden Markov model (HMM) approach using the STAN software (Zacher et al., 2014). These initial TUs were refined by combining disconnected TUs based on the exon-exon junction information in the data from RNA-seq. Next, the TUs were refined to true 5’-ends and 3’-ends based on CAGE and 3’-end-seq data, respectively (Andersson et al., 2014b; Wu et al., 2020). Finally, we overlapped the TUs with the GenCode v21 reference genome to assign the gene names and biotypes for TUs overlapping the reference annotation and determination of maximal region, major TSS and TES and major splice isoforms using customized python scripts.

### 3’-end-seq library construction and data processing

Experiments for 3’-end-seq libraries were conducted in triplicate and the libraries were constructed, sequenced and analyzed as described previously (Wu et al., 2020). However, for this study we only sequenced the 4sU-labeled and -purified samples. The obtained data provided a mix of different stable and transient 3’-ends: (1) ‘co-transcriptional’ 3’-ends residing in the active site of RNAPII, (2) ‘post-transcriptional’ 3’-ends (pA^+^ as well as pA^-^) produced either by RNA processing or directly by transcription termination, and (3) 3’-ends from intermediates of the splicing process ((Wu et al., 2020), and data not shown). Various filtering controls revealed that the mixed nature of the samples did not skew the conclusions of this study (data not shown). Moreover, there were subtle differences between the replicates, but these were irrelevant to the overarching conclusions for this study.

### Identification of INT-sensitive sites

In order to pinpoint 3’-ends that respond to INT depletion, we first selected sites from normalized and genomic A-filtered EXO and EXO/INTS ‘all 3’-ends’ samples (three replicates each) which contained at least 100 normalized reads across the 6 samples, which gave us 58,549 sites for further analysis. Sites were then deemed as INT-sites if they were significantly ‘downregulated’ in the INT/EXO samples compared to EXO samples based on unpaired, two-sided Student t-tests of log2 transformed values after adding a pseudocount of 1, selecting sites with decreased signal and Benjamini-Hochberg corrected p-values < 0.1. The resulting 3,928 individual positions were then used for further analysis (Table S1).

### Selection of TUs for downstream analysis

To exclude TUs that are deregulated due to readthrough transcription from upstream, we included TUs in the downstream analyses set if the signal coming from upstream of the first TSS did not surpass 0.5-fold compared to mean TU-body signal (indicated in Figure S1E-G). For TSS-anchored analyses, the analyzed subset was further reduced based on the same criteria for the specific TSSs analyzed.

TUs were divided into biotype categories based on Gencode annotation and as described under results for previously unannotated TUs (Figure S1E-F, 2D). Only TUs falling within the ‘top10’ biotypes (top9 based on number of TUs within the category + snRNA genes) were kept for further analysis. TUs were divided into multi- and mono-exonic TUs based on the presence or absence, respectively, of splice-junction reads in RNA-seq (Figure 2D).

### Aggregate and ‘genome browser’ plotting

Metagene profiles were aggregated by collecting coverage/signal for the window of interest anchored to TSS, TES, pA-site as indicated. These were then binned into 50 nt bins, and the mean coverage/signal was determined for each bin. The 90% confidence interval for the mean was determined by 50 bootstrap samplings with replacement.

For ‘genome browser’-type plots, the coverage for each of the samples at each individual chromosomal position was calculated using customized python and R scripts. The coverage was normalized using the sample-specific size-factors, corrected as described above and stored in wig files. Wig files were then converted to bigwig files using the UCSC wigToBigWig tool (Kent et al., 2010). bigwig files were visualized using a customized R script.

### Readthrough index, initiation index and splice acceptor index

The log2-transformed difference in readthrough levels (ΔRT) used to sort heatmaps in Figure 2G-H was calculated based on coverage within the region up to 1 kb downstream of the TES in knockdown *vs*. CTRL samples normalized to the coverage within the region up to 500 bp upstream of the TES.

The log2-transformed difference in ‘initiation signal’ (ΔIS) used to sort heatmaps in Figure 3C-D was calculated as the difference in log2-transformed mean coverage between knockdown *vs*. CTRL samples over the first 500 bp of the TU.

The splice acceptor index (ΔSA) used to assess differences in splicing was calculated for each individual splice acceptor as the mean (of two replicates) of the log10-transformed ratio between number of ‘spliced’ reads divided by total number of reads covering the splice acceptor.

### Published datasets

To generate the aggregate metagene profiles of published data we downloaded bigwigs containing the normalized coverage. These were then log2-transformed and collected for the window of interest anchored as indicated, binned into 50-bins. The plots were generated as for data collected in this study, the means and confidence intervals were determined for each of these bins using customized R scripts.

Annotation TSSs: (Andersson et al., 2014b); GEO: GSE58991

Annotation TESs: (Wu et al., 2020); GEO: GSE137612

mNET-seq (Figure S4A): (Nojima et al., 2015); GEO: GSE60358

mNET-seq (Figure S4B): (Schlackow et al., 2017); GEO: GSE81662

Chr-seq (Figure S4C): (Nojima et al., 2015); GEO: GSE60358

GRO-seq (Figure S4D): (Gardini et al., 2014); GEO GSE58255

### Statistical analysis

All statistical analyses are indicated in the legends to the relevant figure.

### Data and Code Availability

The datasets generated during this study are available at GEO accession code: pending.

## SUPPLEMENTAL INFORMATION TITLES AND LEGENDS

**Table S1 | INT-sensitive sites**

**Figure S1.**
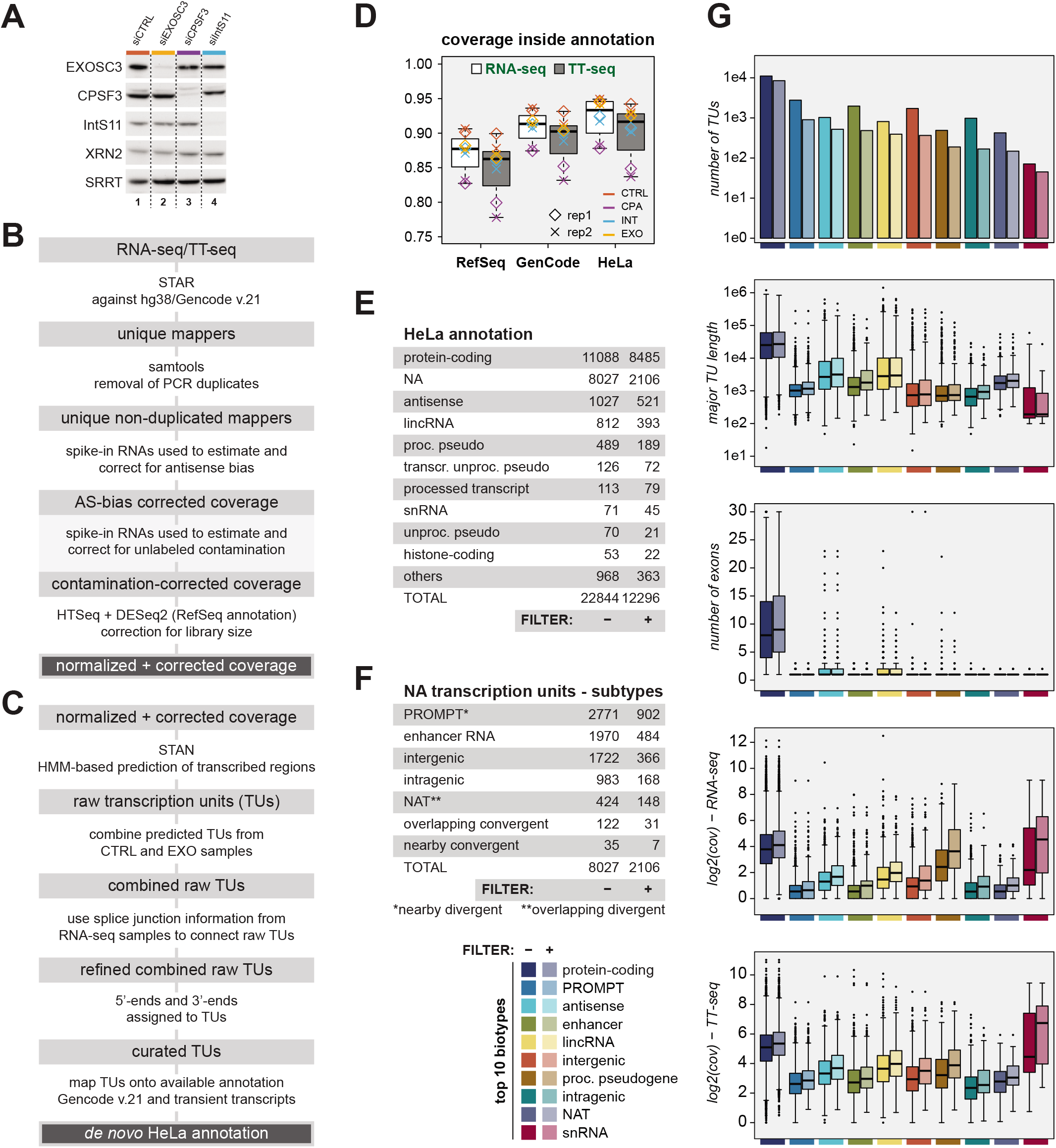
Exhaustive *de novo* annotation of HeLa cell TUs (related to Figure 1) (A) Western blotting analysis demonstrating efficient knockdown of indicated proteins. XRN2 and SRRT/ARS2 were probed as loading controls. (B) Flow diagram illustrating the computational workflow for preprocessing the TT-seq and RNA-seq data to obtain coverage corrected for antisense-bias, contaminating unlabeled RNA (in the case of TT-seq) and library-size. (C) Flow diagram illustrating the computational steps applied to TT-seq and RNA-seq data from siEGFP- and siEXOSC3-treated HeLa cells to annotate the HeLa cell transcriptome. (D) Box plot displaying the fraction of RNA-seq (white) and TT-seq (gray) signal coverage from the indicated samples (siEGFP-, siCPSF3-, siIntS11- and siEXOSC3-treated samples, coined CTRL, CPA, INT and EXO, respectively), falling inside RefSeq, Gencode or *de novo* HeLa annotations. Box limits represent the first and third quartiles, the band inside the box is the median. The ends of the whiskers extend the box by 1.5 times the interquartile range. (E) Overview of biotypes present in the HeLa transcriptome. NA=not annotated in Gencode v.21. The ‘FILTER column displays the number of identified TUs. The ‘FILTER +’ column displays the number of TUs used for downstream analysis in this study, which did not have substantial signal entering from the upstream locus in any of the analyzed samples (see *Materials and Methods* for details). (F) Overview of subtypes of NA TUs. Columns as in (E). (G) Characteristics of the TUs within the major (top 10, color-coded) biotype-classes represented in the HeLa transcriptome annotation. For each biotype the values are shown for the complete annotated set and the subset used in downstream analyses on the left and right, respectively. From the top: (1) number of TUs within biotype-class (log10-scale), (2) length of TU (log10-scale), (3) the number of exons, (4) median log2-transformed RNA-seq coverage and (5) median log2-transformed TT-seq coverage. For clarity, the y-axis in (1) was restricted at 30 exons although there are several protein-coding TUs with more exons. Values are shown for the major transcript (see *Materials and Methods* for details). Median, hinges and whiskers in box plots (panels 2-5) are shown as in (D).

**Figure S2.**
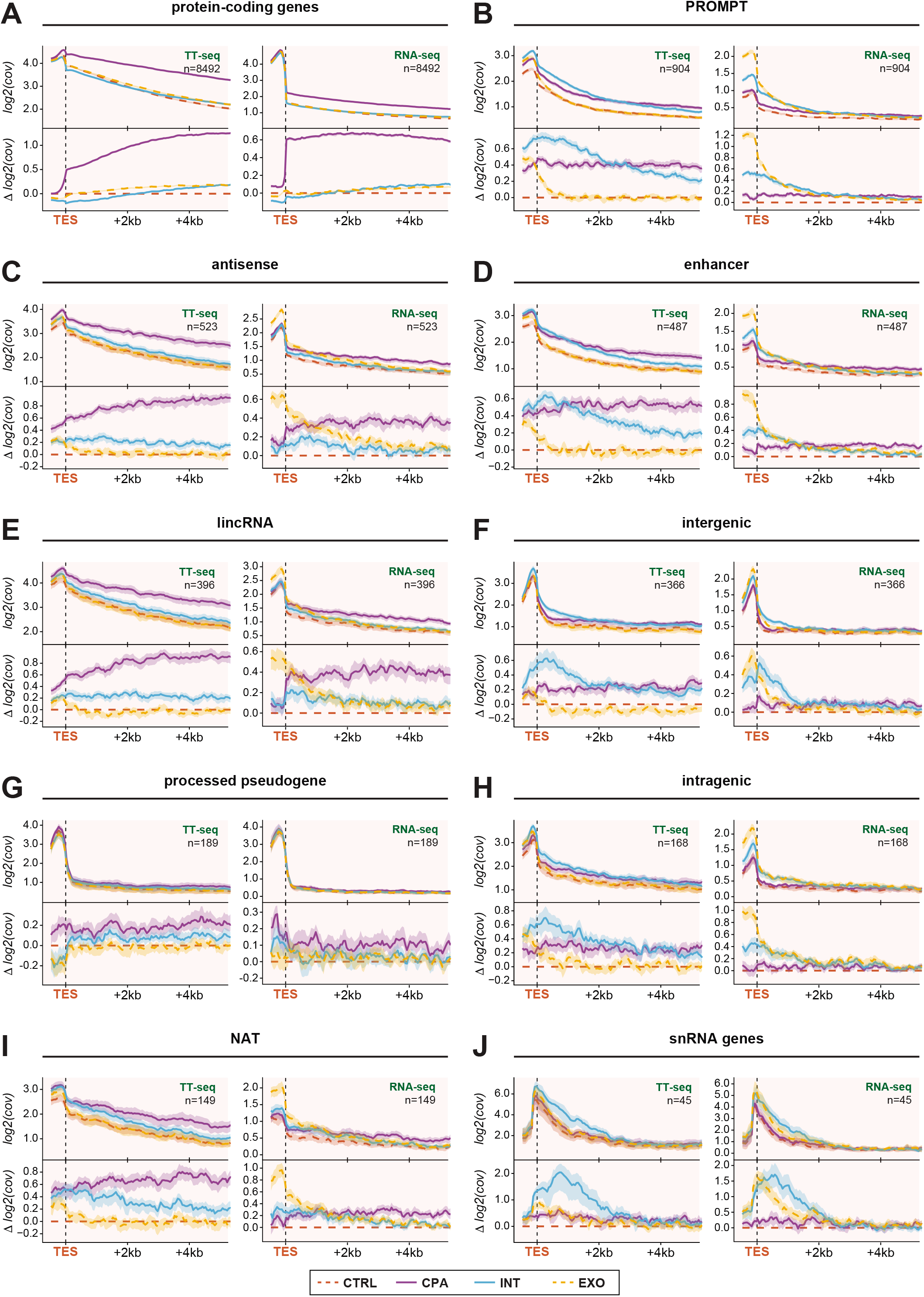
CPA and INT depletions display common and diverse global transcription termination phenotypes at TU ends (related to Figure 2) (A)-(J) Aggregate plots of mean TT-seq (*left*) and RNA-seq (*right*) data aligned to TESs of TUs from major RNA biotype-classes represented in the HeLa transcriptome annotation: (A) protein-coding, (B) PROMPT, (C), antisense, (D) enhancer, (E) lincRNA, (F) intergenic, (G) processed pseudogene, (H) intragenic, (I) NAT and (J) snRNA. Representation as in Figure 2E.

**Figure S3.**
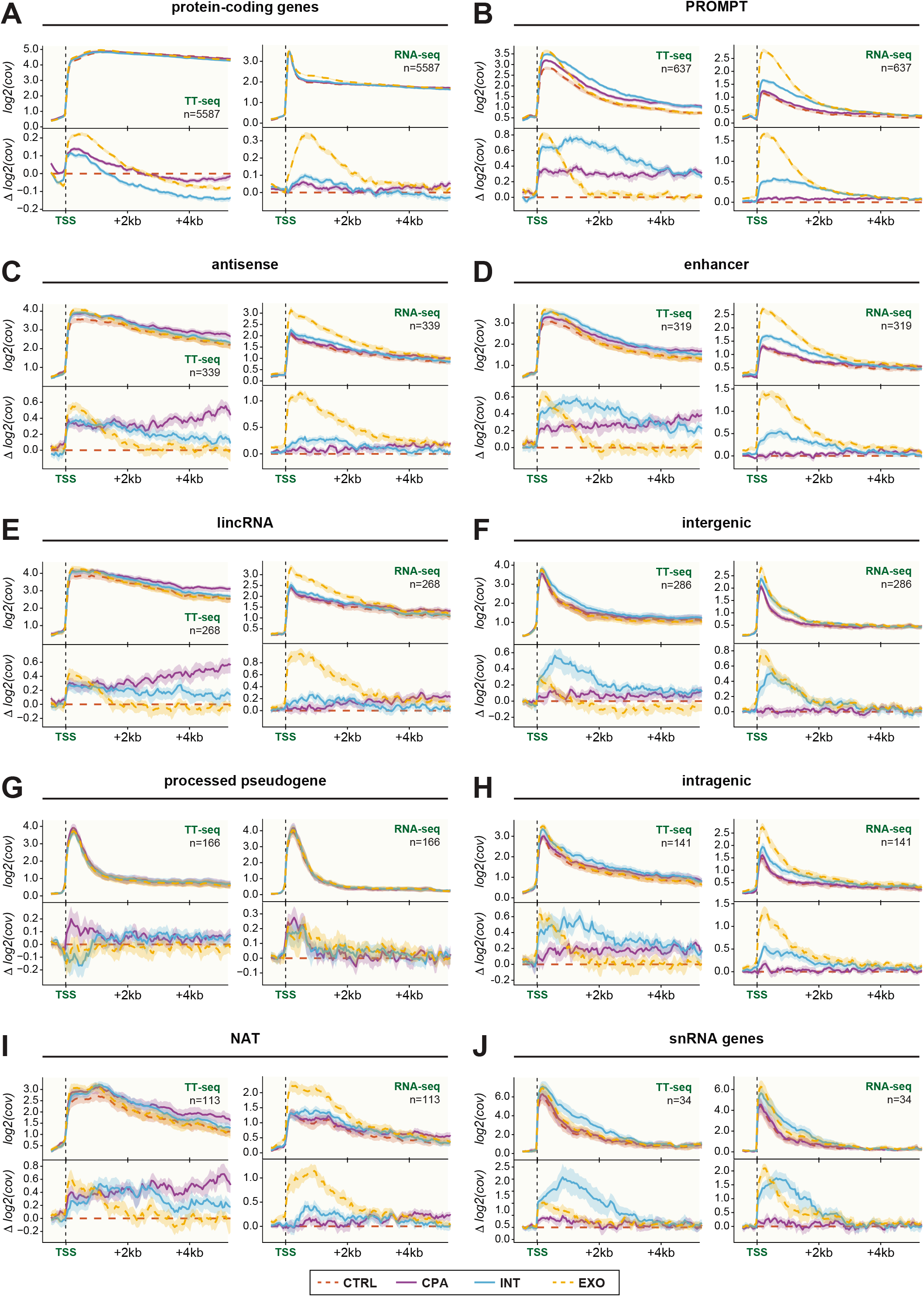
CPA and INT depletions cause increased promoter-proximal transcription globally (related to Figure 3) (A)-(J) Aggregate plots of mean TT-seq (*left*) and RNA-seq (*right*) data aligned to TSSs within TUs from major RNA biotype-classes represented in the HeLa transcriptome annotation: (A) protein-coding, (B) PROMPT, (C), antisense, (D) enhancer, (E) lincRNA, (F) intergenic, (G) processed pseudogene, (H) intragenic, (I) NAT and (J) snRNA. Representations as in Figure 3A.

**Figure S4.**
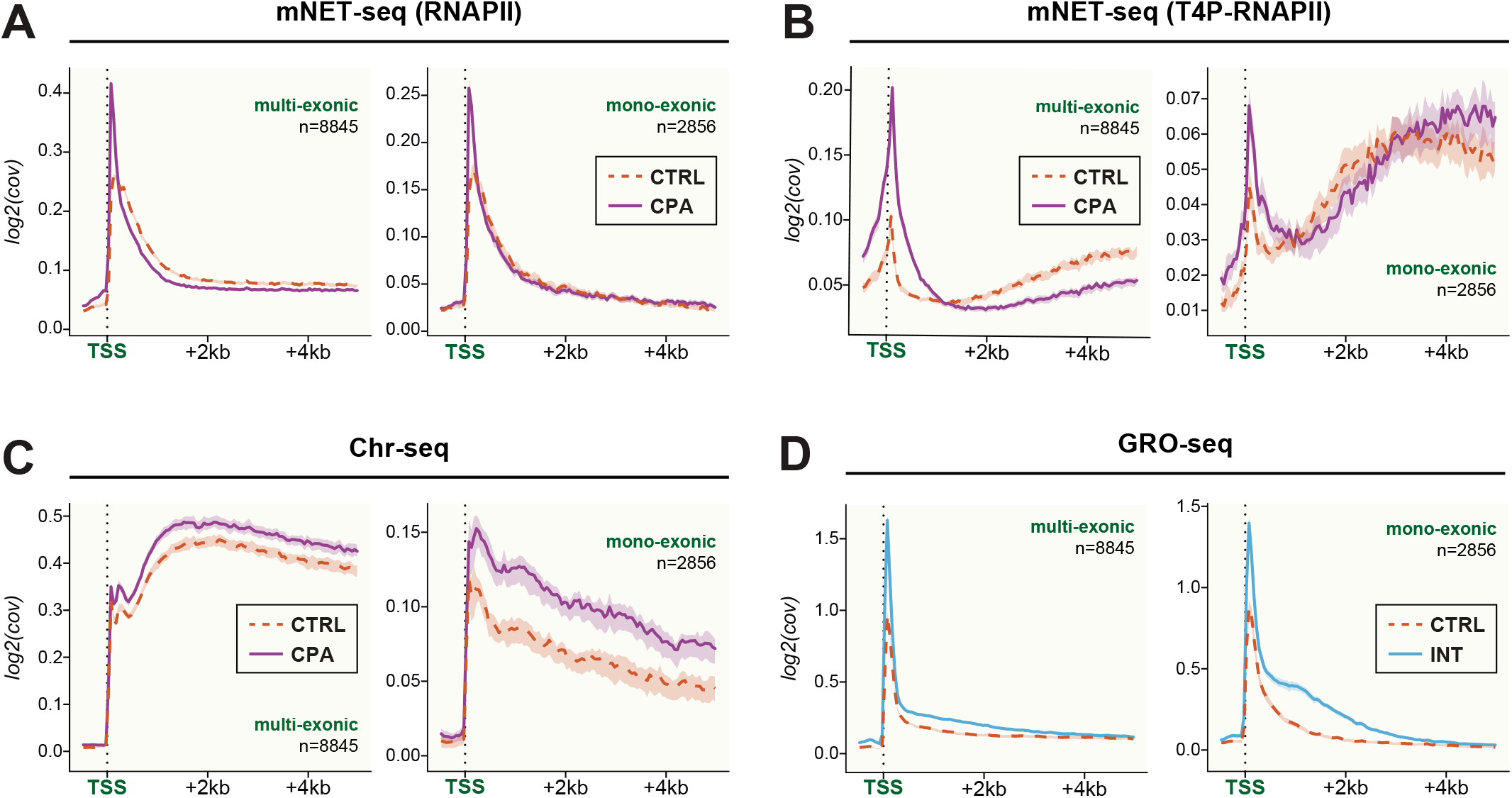
CPA and INT depletions cause increased promoter-proximal transcription globally (related to Figure 3) (A) Aggregate plots of mean total mNET-seq data for CPA depletion (*purple*) and CTRL (*red*) aligned at the TSS of multi-exonic (*left*) and mono-exonic (*right*) TUs. Mean values and 90% confidence intervals of log2-transformed coverage for 50-bp bins over all included TUs are displayed. The number of aggregated loci (n) is indicated. (B) Aggregate plots of mean T4P-RNAPII mNET-seq data for CPA depletion and CTRL aligned at the TSS of multi-exonic (*left*) and mono-exonic (*right*) TUs. Representations as in (A). (C) Aggregate plots of mean Chr-seq data for CPA depletion and CTRL aligned at the TSS of multi-exonic (*left*) and mono-exonic (*right*) TUs. Representations as in (A). (D) Aggregate plots of mean GRO-seq data for INT depletion (*cyan*) and CTRL (*red*) aligned at the TSS of multi-exonic (*left*) and mono-exonic (*right*) TUs. Representations as in (A).

**Figure S5.**
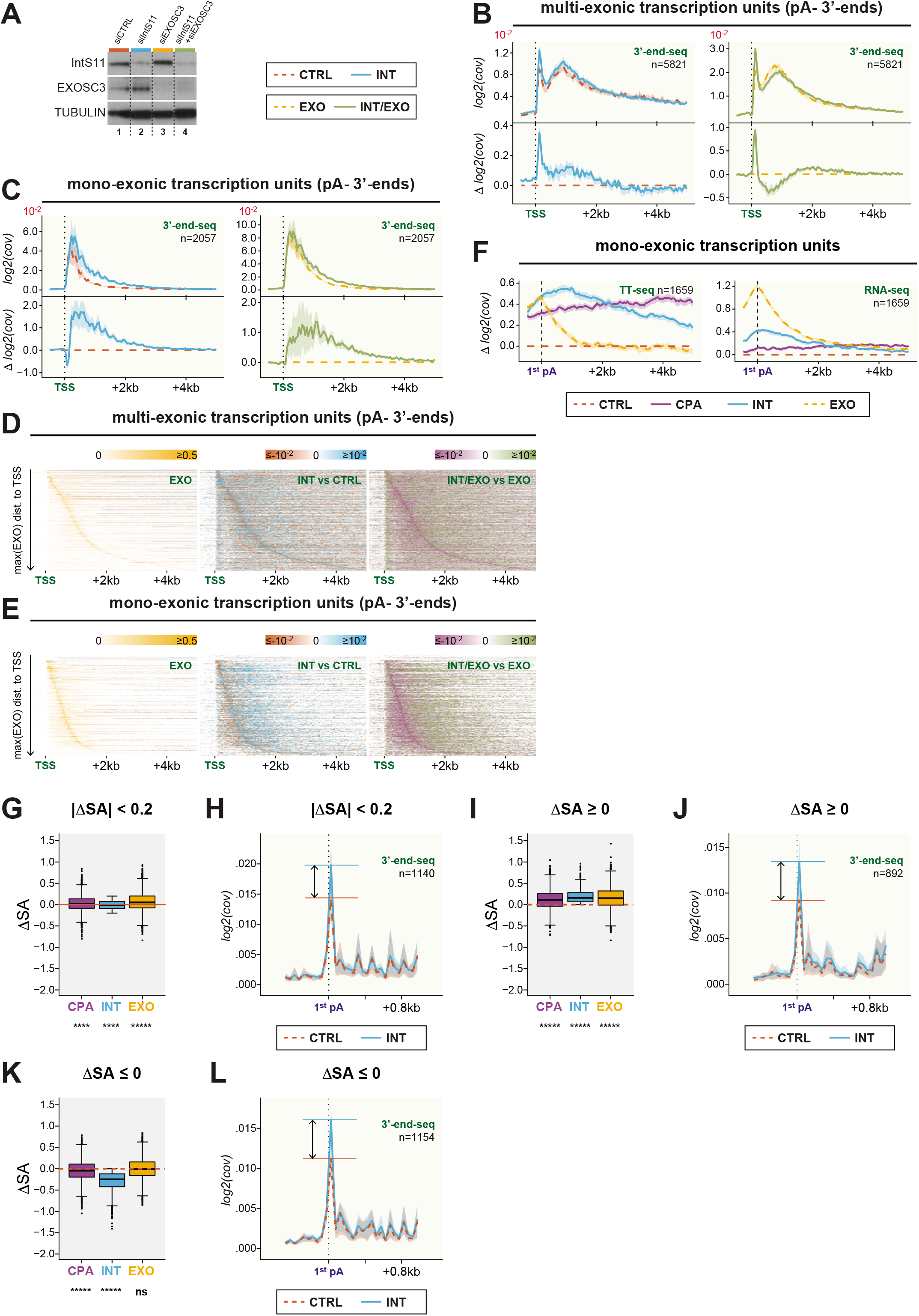
INT depletion exposes non-productive RNAPII transcription events (related to Figure 4 and 5) (A) Western blotting analysis demonstrating efficient knockdown of indicated proteins. Tubulin was probed as loading control. (B)-(C) Aggregate plots as in Figure 4B-C, but of pA^-^ 3’-end-seq data. (D)-(E) Heatmaps as in Figure 4D-E, but of pA^-^ 3’-end-seq data. (F) Aggregate plots as in Figure 5E but for mono-exonic TUs. (G) Box plot displaying the changes in splice acceptor ratios (ΔSA) for CPA (*purple*), INT (*cyan*) and EXO (*orange*) depletions relative to CTRL for the subset of multi-exonic TUs with |ΔSA|≤0.2 for INT depletion samples. Box limits represent the first and third quartiles, the band inside the box is the median. The ends of the whiskers extend the box by 1.5 times the interquartile range. (H) Aggregate plots of pA^+^ 3’-end-seq data aligned at the first encountered putative pA signal (1^st^ pA) of multi-exonic TUs with |ΔSA|≤0.2 for INT depletion (*cyan*) and CTRL (*red*) samples. Mean values and 90% confidence intervals of log2-transformed coverage for 50-bp bins are displayed. Horizontal lines in corresponding colors indicate the peak height and the arrows indicate the difference between the peak height. (I)-(J) and (K)-(L) As (G)-(H), but for a subset of multi-exonic TUs with ΔSA≥0 and ΔSA≤0 for INT depletion, respectively. For the box plots in (G), (I) and (K), the ΔSA distributions were subjected to Wilcoxon tests of significance. *p < 0.05, **p < 0.01, ***p < 0.005, ****p < 0.001, *****p < 2.2e-16, ns, not significant.

**Figure S6.**
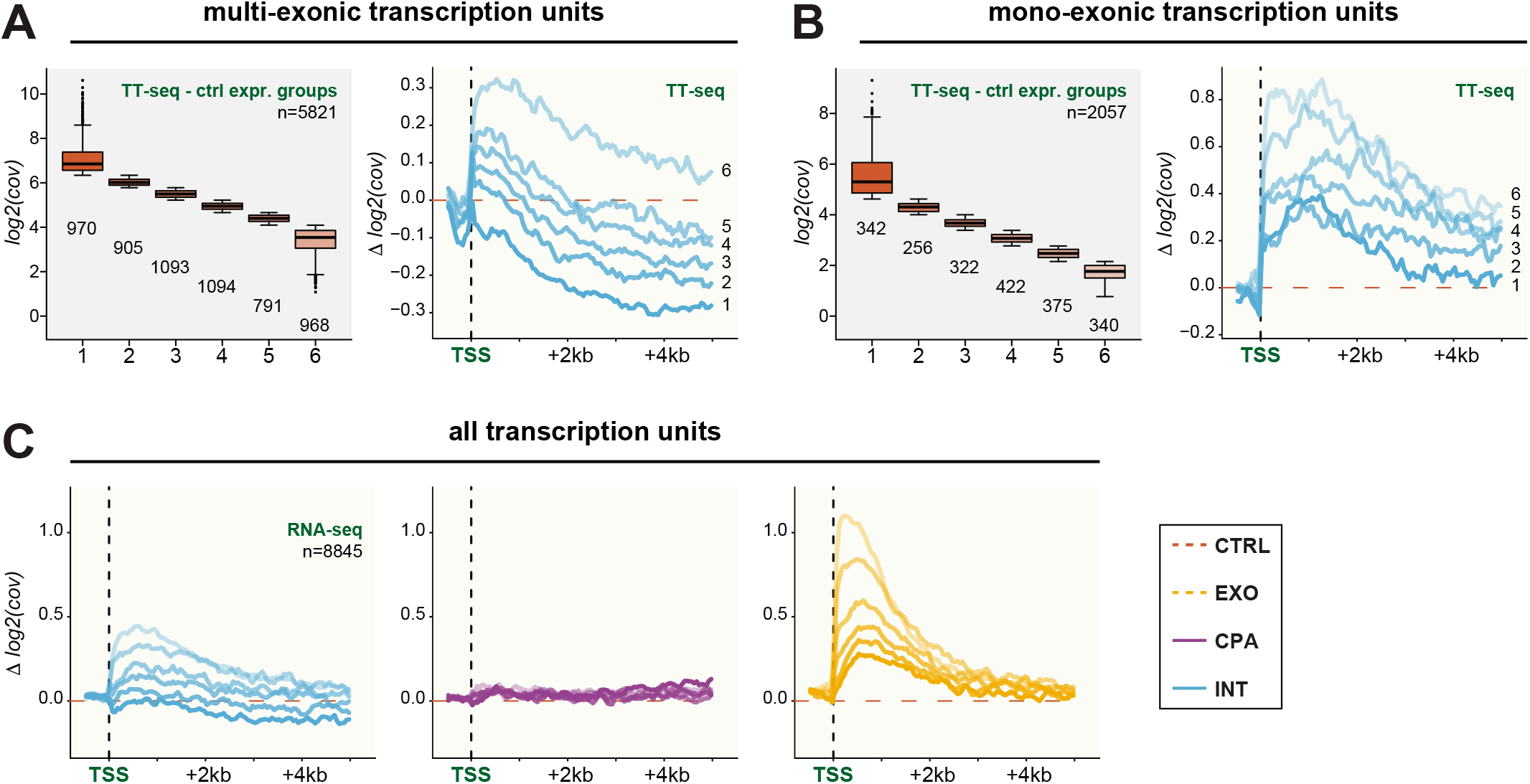
INT activity correlates inversely with transcriptional output. (A) Box plot displaying expression levels for six expression-based multi-exonic TU groups (1-6) based on expression in CTRL (*left*). Box limits represent the first and third quartiles, the band inside the box is the median. The ends of the whiskers extend the box by 1.5 times the interquartile range. Outliers are show in black. The number of TUs in each group is indicated. Aggregate plots (*right*) of the log2-transformed TT-seq Δ coverage of INT (*cyan*) depletion vs. CTRL aligned at the major TSS. Mean values of log2-transformed Δ coverage for 50-bp bins are displayed. The line color intensity (high to low) corresponds to the gene group 1 to 6, respectively. (B) As (A), but for six expression-based mono-exonic TU groups (1-6) based on expression in CTRL (*left*). (C) Aggregate plots as in Figure 6B, but for RNA-seq data.

**Figure S7.**
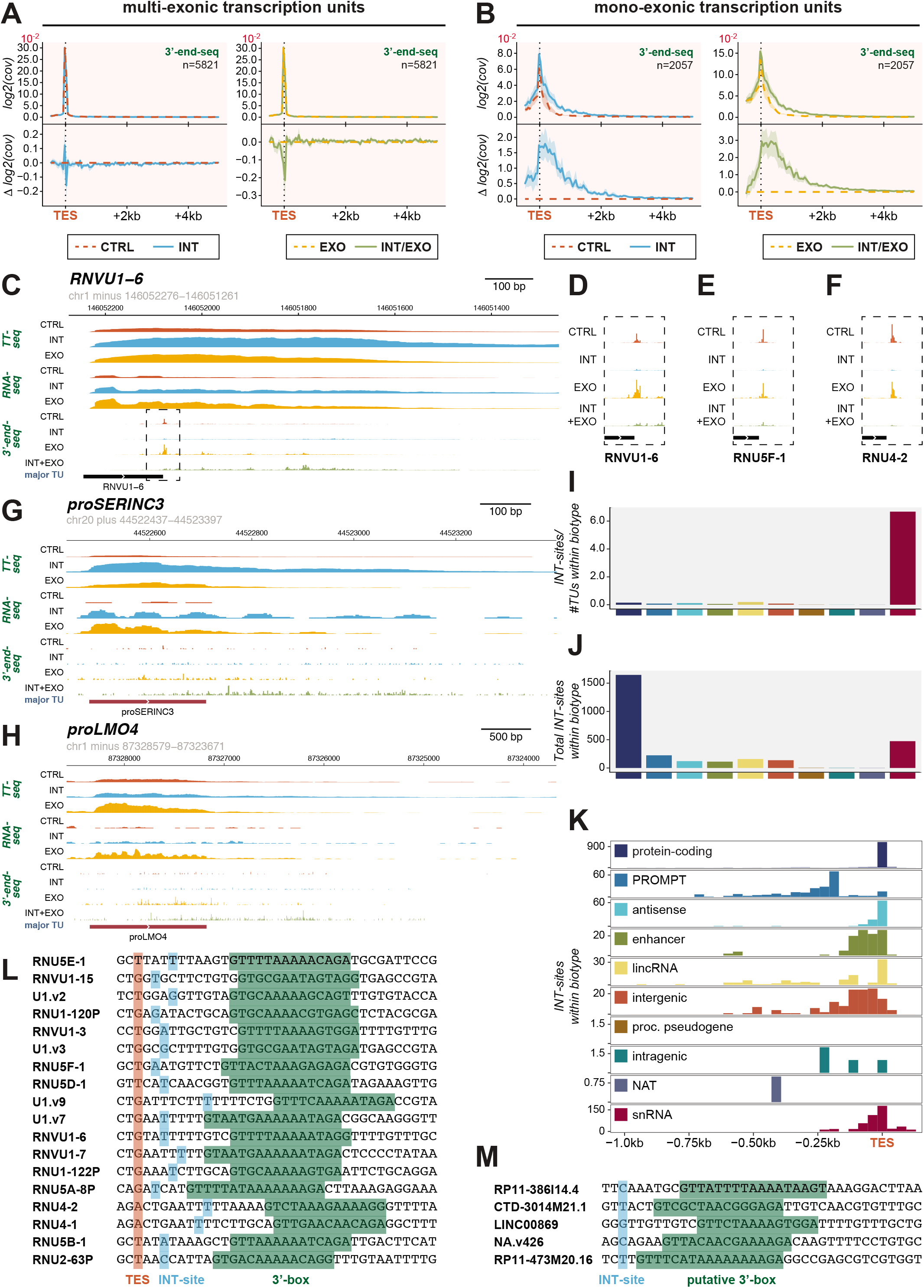
snRNA TUs exploit INT-sensitive transcription for their stable RNA production. (A)-(B) Aggregate plots as in Figure 7A-B, but for multi-exonic and all mono-exonic TUs, respectively. (C) Genome browser view of *RNVU1-6* (snRNA) and downstream region. Mean TT-seq and RNA-seq coverage (of two replicates) on the minus strand from control (CTRL, *red*), INT (*cyan*) and EXO (*yellow*) depleted HeLa cells are shown. Additionally, mean 3’-end-seq data (of three replicates) on the minus strand from control (CTRL, *red*), INT (*cyan*), EXO (*yellow*) and INT/EXO (*olive green*) depleted HeLa cells are shown. Rectangle with dashed outline indicate region that is enlarged in (D). (D) Enlarged plots of the 3’-end-seq data at the 3’-end of *RNVU1-6*. (E)-(F) as in (D), but for the snRNA TUs *RNU5F-1* and *RNU4-2*. (G)-(H) As (C), but for the PROMPT TUs *proSERINC3* and *proLMO4*. (I)-(K) As Figure 7C-E, but for the ‘top10’ biotypes as defined in Figure S1. (L) Sequences from the indicated snRNA TUs aligned to TES (shaded red; positions −2 to +34 are shown). TUs with INT-sensitive site −5 to +10nt from TES are shown. INT-sensitive 3’-end sites (INT-site) are shaded cyan. Putative 3’-boxes (based on (Hernandez, 1985)) are shaded green. Motif emerging from here: INT-sensitive site -N_2-9_GTttN_0-3_AAaADN_1-2_AGR (where upper- and lowercase letters indicate strict and less strict adherence to motif, N is any nucleotide, D=A,G or T, R=A or G). Note the most significant INT-sensitive site is often, but not always the most abundant 3’-end in the region. (M) As (L), but for selected non-snRNA TUs. Sequences are aligned to the INT-sensitive 3’-end site. *RP11-386I14.4* (antisense; also known as *proDNAJB4), CTD-3014M21.1* (TEC), *LINC00869* (lincRNA), *NA.v426* (intergenic) and *RP11-473M20.16* (lincRNA).

